# Exposure to Maternal Pre- and Postnatal Psychological Distress: Associations with Brain Structure in 5-year-old Children

**DOI:** 10.1101/2025.11.24.690168

**Authors:** Elmo P. Pulli, Hilyatushalihah K. Audah, Amira Svensk, Aylin Rosberg, Silja Luotonen, Pauliina Juntunen, Isabella L. C. Mariani Wigley, Venla Kumpulainen, Eero Silver, Anni Copeland, Ekaterina Saukko, Harri Merisaari, Eeva-Leena Kataja, Linnea Karlsson

## Abstract

**Background:** Maternal mental health is an important contributor to child neurodevelopment. While there are multiple studies on prenatal exposure, early postnatal exposure has received little attention in neuroimaging research.

**Methods:** 5-year-old children (n = 173) were recruited from the FinnBrain Birth Cohort study. Maternal distress was assessed using questionnaires on depressive and anxiety symptoms at 14, 24 and 34 gestational weeks and postnatally at 3, 6 and 24 months. T1-weighted structural images were processed using a voxel-based morphometry pipeline to map associations between maternal distress exposure and regional gray matter (GM) volumes, while accounting for potential confounders.

**Results:** We found widespread associations between maternal distress symptoms and offspring brain morphology. Higher prenatal distress at 14 gestational weeks was positively associated with regional GM volume in the right superior parietal lobe and precuneus. In contrast, postnatal distress at 3 months was negatively associated with GM volumes in multiple motor regions, the left anterior insula, right superior frontal areas and supramarginal gyrus. Postnatal distress at 6 months demonstrated a positive relationship with GM volumes in the right calcarine and lingual gyri, while distress at 24 months was negatively associated with GM volumes in the left supramarginal and right superior frontal gyri.

**Conclusions:** This study provides support for hypotheses proposing that fetal and early life exposure to maternal distress can influence the structural development of the brain. Furthermore, it highlights the role of early postnatal period and calls for further research into this so far overlooked period and pathways that explain the associations.

## Introduction

Maternal pre- and postnatal mental distress (e.g., depressive or anxiety symptoms), may present significant, enduring, and potentially irreversible effects on offspring development [1–4]. Both pre- and postnatal distress share similarities in their potential long-term outcomes [5] which could reflect increased genetic vulnerability [6,7]. It is still useful to study the timing of exposure, as pre- and postnatal distress exposure involve different mechanisms. Prenatal distress can affect fetal development via biological pathways such as perturbation of the hypothalamic–pituitary–adrenal axis function, increased inflammation [8], and epigenetic modifications of the placenta [9], while postnatal maternal distress can affect the offspring through psychosocial and environmental factors, such as maternal sensitivity [10] and attachment patterns [11]. The environment can affect the brain through neural mechanisms such as activation-dependent plasticity, mediated by brain-derived neurotrophic factor [12–14]. As some of the mediating mechanisms are different, pre- and postnatal exposures could have different effects on offspring structural brain development, which can be studied using magnetic resonance imaging (MRI).

Prior neuroimaging research has put much focus on maternal prenatal as opposed to postnatal distress. Common measures include maternal depressive, anxiety, and stress symptoms based on questionnaires [2]. Furthermore, structural neuroimaging studies have typically focused on subcortical structures [15–18], especially the amygdala [16,19–27] and the hippocampus [16,24,28]. Previous amygdala studies have found positive [21,29] and negative [16,25,26] associations between prenatal maternal distress and amygdala volume as well as different association based on child sex [21–23], nonlinear relationships [15], or no associations at all [24,30]. Hippocampal findings have been similarly inconsistent. Some studies reported smaller left [24], smaller right [31], and larger bilateral [29] hippocampal volume in newborns associated with maternal prenatal stress. Other studies have focused on gene–environment interactions influencing right hippocampal volume [28,32], while other studies found no significant hippocampal volume differences [16,22,30]. Prenatal distress has also been associated with white matter structure [33] and brain function in the offspring [34].

Associations between prenatal distress exposure and structural cortical brain development in children have been examined in a few studies, all of them showing significant associations in multiple brain regions [35–39]. Wei et al. [35] found that prenatal maternal depressive symptoms associated positively with areas in the prefrontal cortex, superior temporal gyrus, and superior parietal lobe in 4–6-year-old males (opposite pattern in females). El Marroun et al. [36] found positive associations between prenatal maternal depressive symptoms and offspring surface area in the left caudal middle frontal, left superior frontal, and left lateral occipital regions in 6–10-year-olds. Additionally, they found a negative association between prenatal maternal depressive symptoms and cortical thickness (CT) in the left superior frontal gyrus. Davis et al. [37] found an association between exposure to maternal perceived stress and morphology in multiple frontal and temporal regions. Sandman et al. [38] found negative associations between maternal prenatal depressive symptoms and CT in widespread clusters across the cortex in 6–9-year-olds. Buss et al. [39] found associations between pregnancy-related anxiety at 19 gestational weeks (GW) and reductions in gray matter (GM) volumes in regions including the prefrontal cortex, premotor cortex, lateral temporal cortex, medial temporal lobe, postcentral gyrus, and the cerebellum. Although cortical imaging has received less attention than subcortical, there are some regions that are found repeatedly, including frontal and temporal regions.

Postnatal maternal mental health has received less attention in neuroimaging studies, although it has adverse effects on child development comparable to prenatal exposure [5]. Two studies in 2– 3-year-olds found no associations between postnatal depression exposure and CT or area [36,40], while one of them did find smaller amygdala volumes in those exposed to postnatal depression [40]. In older children, exposure to maternal depression has been associated with: smaller right superior frontal CT in 2.6–5.1-year-olds [41], smaller fusiform gyrus volume in 7–15-year-olds [42], and total GM volume in 10-year-olds [43]. Overall, the cortical structural neuroimaging literature on the associations of early life postnatal maternal distress exposure is limited and the results are limited to individual ROIs that do not replicate between studies.

The aim of this study was to investigate the associations between pre- and postnatal maternal stress and brain structure in 5-year-old children. Distress was measured longitudinally utilizing questionnaire data on depressive and anxiety symptoms from GWs 14, 24, and 34 as well as from 3, 6, and 24 months of child age postnatally. We expected to find: 1) significant differences in GM volumes in fronto–temporal regions; and 2) alterations in amygdalar and hippocampal volumes in those exposed to pre- and/or postnatal stress. As previous studies combining pre- and postnatal distress symptoms are scarce, we reported exploratory whole-brain VBM analyses.

## Methods and Materials

This study adheres to the ethical guidelines in the 1975 Declaration of Helsinki and has been approved by the joint Ethics Committee of the University of Turku and the Hospital District of Southwest Finland (ETMK:31/180/2011). This study follows the Strengthening the Reporting of Observational Studies in Epidemiology (STROBE) guidelines [44], checklist in Table S1.

### Participants

The participants of the present study are mother–child dyads (n = 173) drawn from the FinnBrain Birth Cohort study (www.finnbrain.fi)[45]. Children who underwent neuroimaging during their 5-year follow-up visit (n = 203, 55.7% boys, mean age 5.39 [SD 0.13], range 5.08–5.79 years) were eligible to participate. Exclusion criteria for the MRI included the following: premature birth (< 35 GW, or < 32 GW for those with exposure to maternal prenatal synthetic glucocorticoid treatment); developmental sensory anomaly or disability (e.g., blindness); long-term medical conditions (e.g., epilepsy, autism spectrum disorders); current ongoing medical examinations or clinical follow-ups; usage of daily medications; history of head trauma; routine MRI contraindications. More details on the cohort and recruitment process in Supplementary Materials. Participant characteristics are presented in Table 1.

**Table 1.**
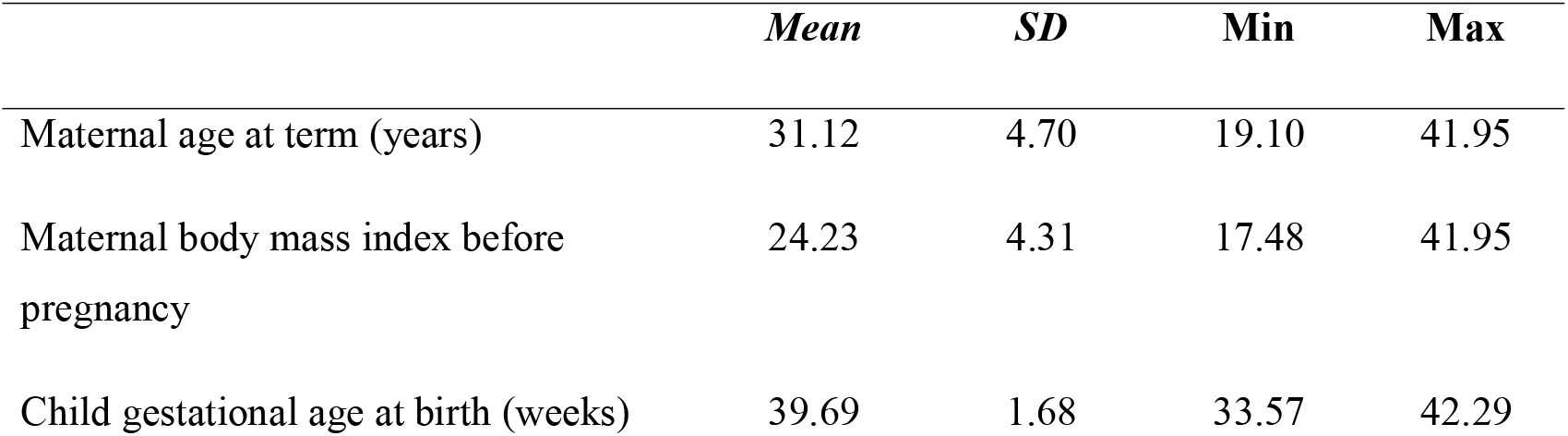
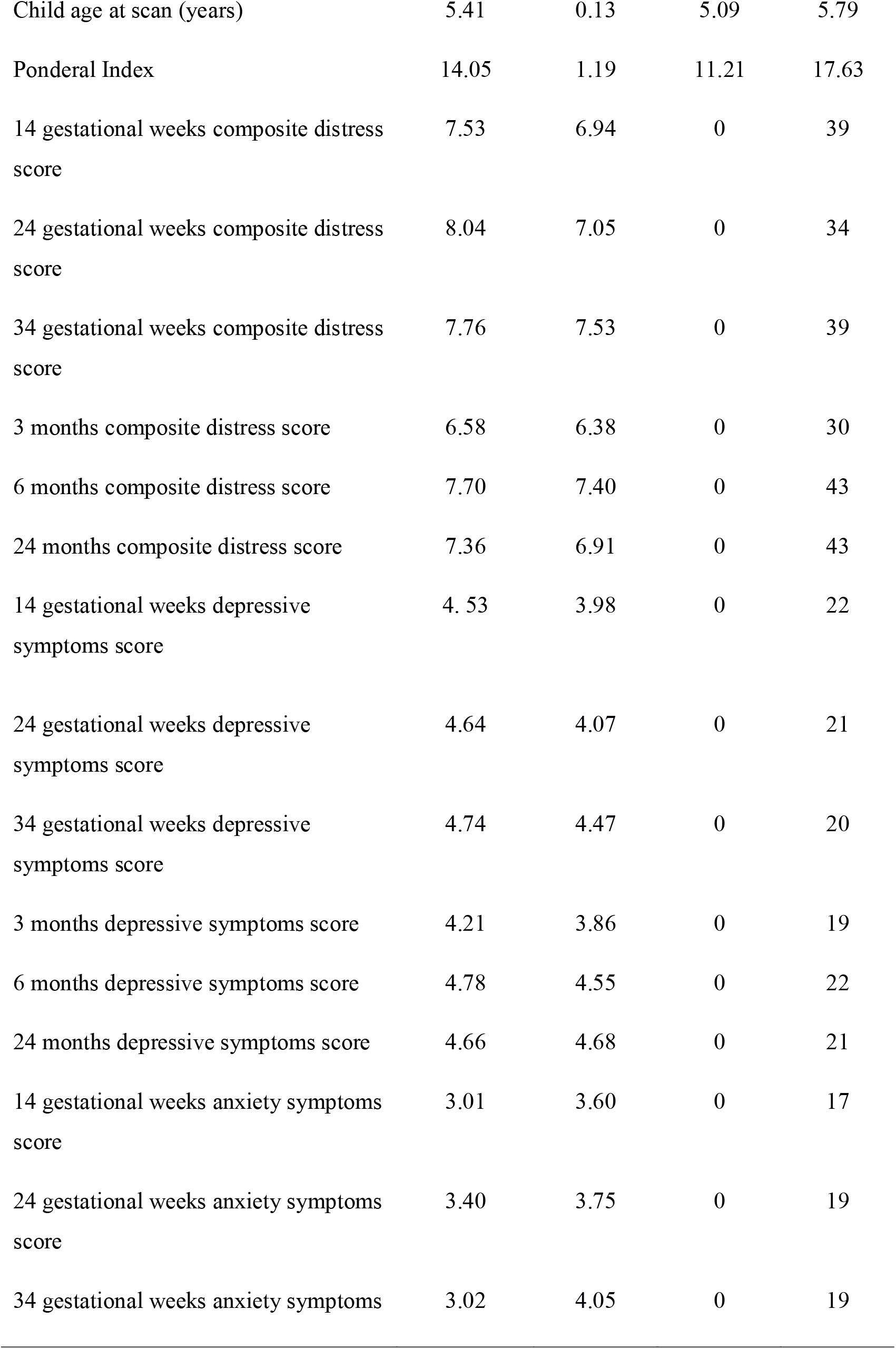
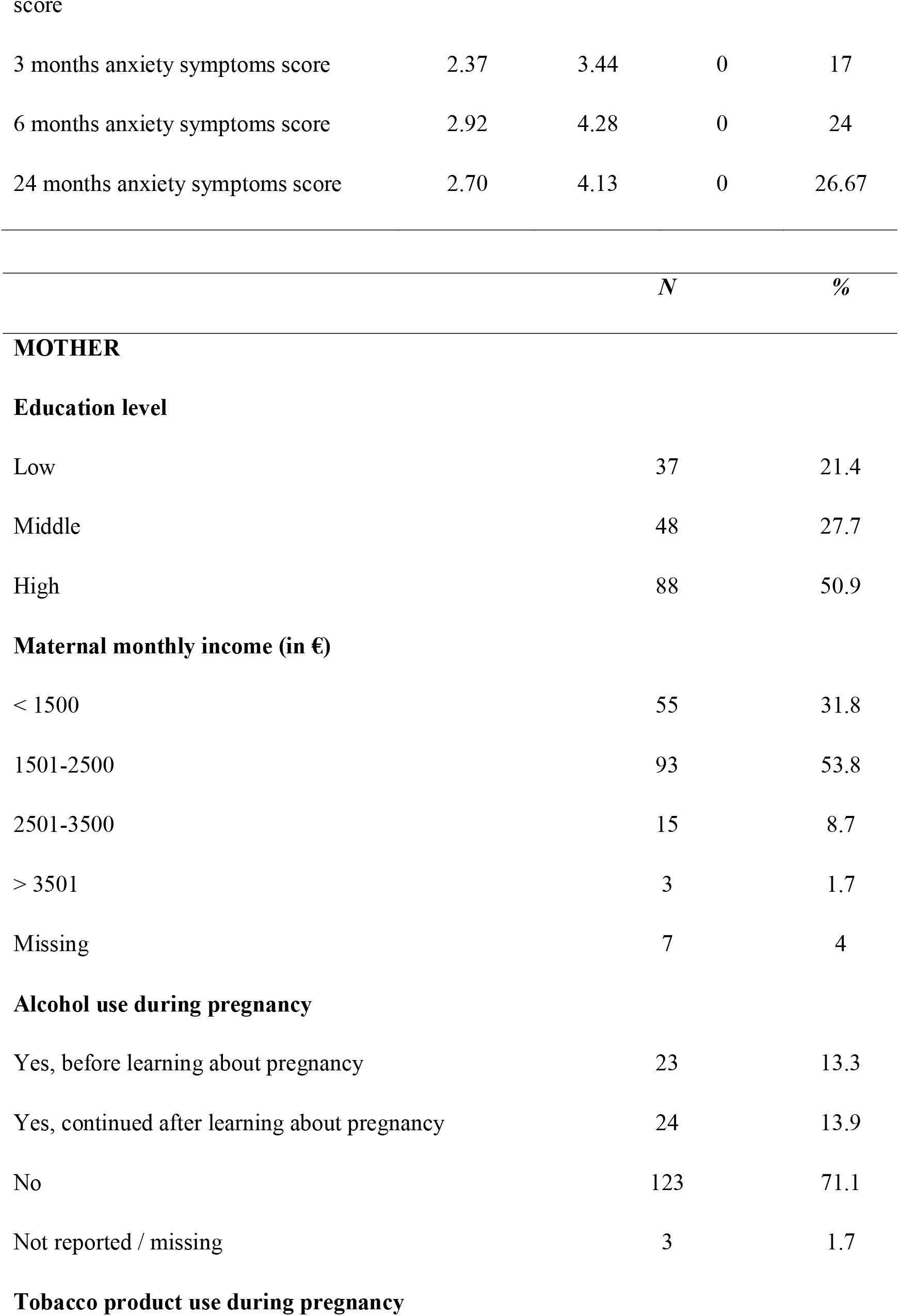
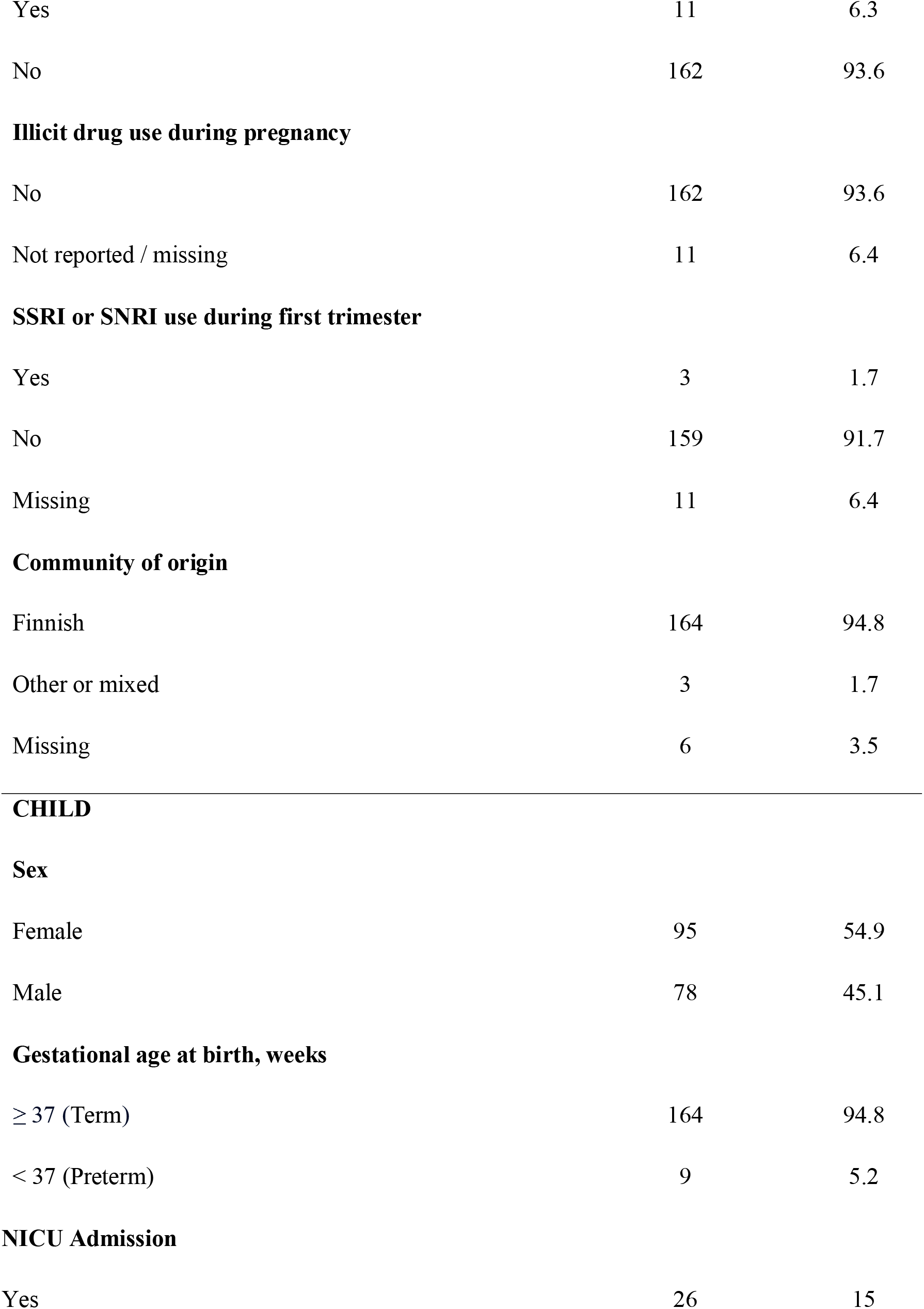
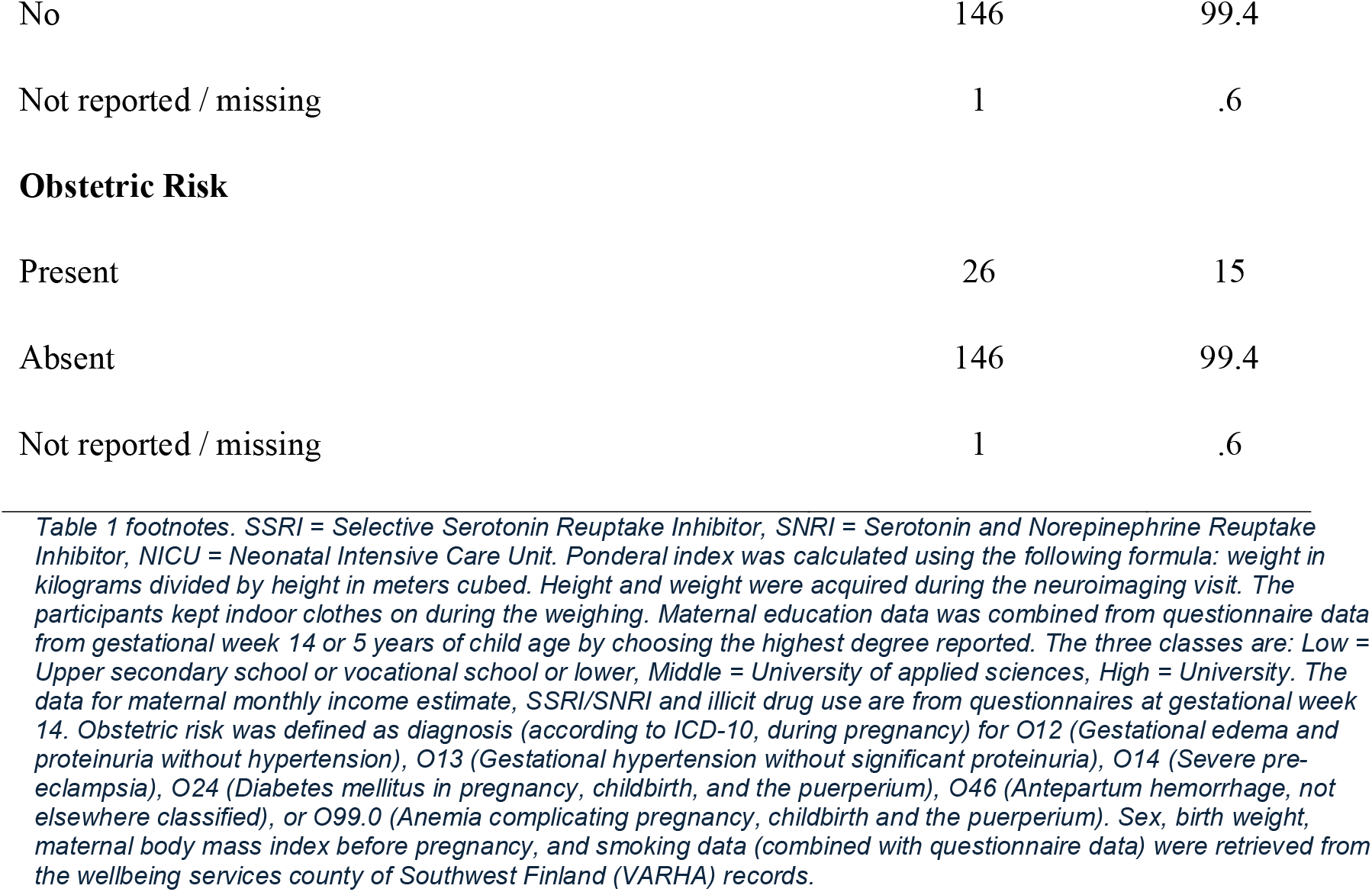
Demographic variables and distress scores of participating mothers and children.

|  | *Mean* | *SD* | **Min** | **Max** |
| --- | --- | --- | --- | --- |
| Maternal age at term (years) | 31.12 | 4.70 | 19.10 | 41.95 |
| Maternal body mass index before pregnancy | 24.23 | 4.31 | 17.48 | 41.95 |
| Child gestational age at birth (weeks) | 39.69 | 1.68 | 33.57 | 42.29 |
| Child age at scan (years) | 5.41 | 0.13 | 5.09 | 5.79 |
| Ponderal Index | 14.05 | 1.19 | 11.21 | 17.63 |
| 14 gestational weeks composite distress score | 7.53 | 6.94 | 0 | 39 |
| 24 gestational weeks composite distress score | 8.04 | 7.05 | 0 | 34 |
| 34 gestational weeks composite distress score | 7.76 | 7.53 | 0 | 39 |
| 3 months composite distress score | 6.58 | 6.38 | 0 | 30 |
| 6 months composite distress score | 7.70 | 7.40 | 0 | 43 |
| 24 months composite distress score | 7.36 | 6.91 | 0 | 43 |
| 14 gestational weeks depressive symptoms score | 4.53 | 3.98 | 0 | 22 |
| 24 gestational weeks depressive symptoms score | 4.64 | 4.07 | 0 | 21 |
| 34 gestational weeks depressive symptoms score | 4.74 | 4.47 | 0 | 20 |
| 3 months depressive symptoms score | 4.21 | 3.86 | 0 | 19 |
| 6 months depressive symptoms score | 4.78 | 4.55 | 0 | 22 |
| 24 months depressive symptoms score | 4.66 | 4.68 | 0 | 21 |
| 14 gestational weeks anxiety symptoms score | 3.01 | 3.60 | 0 | 17 |
| 24 gestational weeks anxiety symptoms score | 3.40 | 3.75 | 0 | 19 |
| 34 gestational weeks anxiety symptoms | 3.02 | 4.05 | 0 | 19 |
| score |  |  |  |  |
| 3 months anxiety symptoms score | 2.37 | 3.44 | 0 | 17 |
| 6 months anxiety symptoms score | 2.92 | 4.28 | 0 | 24 |
| 24 months anxiety symptoms score | 2.70 | 4.13 | 0 | 26.67 |
|  |  |  | *N* | *%* |

**Education level**
|  |  |  |
| --- | --- | --- |
| Low | 37 | 21.4 |
| Middle | 48 | 27.7 |
| High | 88 | 50.9 |

**Maternal monthly income (in €)**
|  |  |  |
| --- | --- | --- |
| < 1500 | 55 | 31.8 |
| 1501-2500 | 93 | 53.8 |
| 2501-3500 | 15 | 8.7 |
| > 3501 | 3 | 1.7 |
| Missing | 7 | 4 |

**Alcohol use during pregnancy**
|  |  |  |
| --- | --- | --- |
| Yes, before learning about pregnancy | 23 | 13.3 |
| Yes, continued after learning about pregnancy | 24 | 13.9 |
| No | 123 | 71.1 |
| Not reported / missing | 3 | 1.7 |

**Tobacco product use during pregnancy**
|  |  |  |
| --- | --- | --- |
| Yes | 11 | 6.3 |
| No | 162 | 93.6 |

**Illicit drug use during pregnancy**
|  |  |  |
| --- | --- | --- |
| No | 162 | 93.6 |
| Not reported / missing | 11 | 6.4 |

**SSRI or SNRI use during first trimester**
|  |  |  |
| --- | --- | --- |
| Yes | 3 | 1.7 |
| No | 159 | 91.7 |
| Missing | 11 | 6.4 |

**Community of origin**
|  |  |  |
| --- | --- | --- |
| Finnish | 164 | 94.8 |
| Other or mixed | 3 | 1.7 |
| Missing | 6 | 3.5 |

|  |  |  |
| --- | --- | --- |
| Female | 95 | 54.9 |
| Male | 78 | 45.1 |

**Gestational age at birth, weeks**
|  |  |  |
| --- | --- | --- |
| ≥ 37 (Term) | 164 | 94.8 |
| < 37 (Preterm) | 9 | 5.2 |

**NICU Admission**
|  |  |  |
| --- | --- | --- |
| Yes | 26 | 15 |
| No | 146 | 99.4 |
| Not reported / missing | 1 | .6 |
| **Obstetric Risk** |  |  |
| Present | 26 | 15 |
| Absent | 146 | 99.4 |
| Not reported / missing | 1 | .6 |

### Procedures

Following recruitment, participating mothers provided demographic information and completed a set of standardized and internationally validated self-report questionnaires regarding depressive and anxiety symptoms three times during pregnancy (14, 24 and 34 GW), and three times in the postnatal period (3, 6 and 24 months). MRI data was collected at age 5 years.

### Measures

Maternal depressive symptoms were assessed using the Finnish version of the Edinburgh□Postnatal□Depression□Scale (EPDS)[46]. Maternal anxiety symptoms were evaluated using the Finnish version of the Symptom Checklist-90-Revised (SCL)[47]. For the purposes of this study, only the anxiety subscale was utilized. The 10-item scale assesses signs of nervousness, tension, and feelings of panic. The questionnaires are described in more detail in Supplementary Materials.

Scores from both the EPDS and SCL were added together to create a unified measure of maternal psychological distress, the composite distress score (CDS). The rationale behind creating a combined, overall measure of distress is twofold: (1) study population were drawn from the general population and consequently, individual scores were relatively low, and (2) symptoms of anxiety and depression are often comorbid [48,49]. A correlation matrix of EPDS and SCL scores is presented in Table S2. Thus, combining the measurements may provide a more robust tool to measure overall distress. Previous authors have used composite scores [22,50,51] or principal component approaches [52] in attempt to grasp the overlap of various distress measures. The EPDS and SCL scores were not standardized while calculating the CDS, as that would make the effect sizes not comparable with other studies. For transparency, we additionally report the results of main analyses with the standardized scores.

There were missing maternal questionnaire data at various follow-up points (14 GW n = 9, 24 GW n = 8, 34 GW n = 8; 3 months n = 17, 6 months n = 32, 24 months n = 47), mean imputation of overall sum scores was used to replace the missing data in such cases (mean of the final sample used in analyses). With missing covariate data, mean imputation was used for maternal pre-pregnancy body mass index (one missing case) and mode imputation with categorical variables (Table 1). Missing data is visualized in Figure S1.

### Image acquisition

The neuroimaging visit is described in our earlier publications [53,54] and in the Supplementary Materials.

Participants were scanned on a 3T MRI system (MAGNETOM Skyra fit; Siemens Medical Solutions, Erlangen, Germany) equipped with a 20-channel head/neck coil. To speed up the acquisition process, the Generalized Auto-calibrating Partially Parallel Acquisition technique was employed, with a parallel acquisition technique factor of two applied to all image acquisitions. High-resolution 3D T1-weighted images using the magnetization prepared rapid gradient echo technique were acquired. This acquisition used the following sequence parameters: repetition time = 1,900 ms, echo time = 3.26 ms, inversion time = 900 ms, flip angle = 9 degrees, voxel size = 1.0 × 1.0 × 1.0 mm^3^, and field of view 256 × 256 mm^2^.

### Data preprocessing and statistical analyses

#### Voxel-based morphometry

VBM is a method that enables researchers to study regional GM volume differences between various groups of participants on a voxel-by-voxel basis. In our study, the Statistical Parametric Mapping (SPM12; http://www.fil.ion.ucl.ac.uk/spm/software/spm12/) software running on MATLAB R2016b (MathWorks, MA) was used to identify alterations in regional GM volumes associated with maternal pre- and postnatal distress.

Preprocessing of the data was done with the Computational Anatomy Toolbox (CAT12, version r1363)[55]. A standard preprocessing pipeline, encompassing normalization, segmentation, modulation, quality assessment, and smoothing, was applied with the following specifications: the default tissue probability map provided in SPM was used, affine regularization was done with the European brains template, inhomogeneity correction was medium, affine preprocessing option was “rough”, local adaptive segmentation was performed at medium, skull-stripping was done via the Graph cuts (GCUT) approach, voxel size for normalized images was set to 1.5 mm and internal resampling for preprocessing was fixed at 1.0 mm. Dartel was used for spatial registration and the MNI152 template was used for both Dartel and Shooting templates.

#### Statistics

Correlation analyses performed using IBM SPSS for Mac (Version 29.0; IBM Corporation), and stepwise regression models with Akaike information criterion for region of interest (ROI) data were performed using RStudio (Version 2022.07.1+554)[56]. All analyses were performed two-tailed.

We performed whole-brain general linear models in SPM12 with the CDS at 14, 24, and 34 GW and at 3, 6, and 24 months postpartum as covariates of interest. Based on findings of previous FinnBrain publications on the same age group [57], the following covariates were included in analyses: child age at scan, sex, and ponderal index, maternal age at term and education level (two classes: (1) any university degree, and (2) other degrees). Analyses were performed with a voxel-wise threshold of p < 0.005 uncorrected. However, we only report clusters that remained significant (p < 0.05) after correction for multiple comparisons using FDR approach. In addition to CDS, we performed the same VBM analyses for EPDS scores to test whether the results of exposure to depressive symptoms, specifically, differ from exposure to distress (with same voxel and cluster thresholds).

The MarsBaR toolbox [58] within SPM12 was used to extract significant clusters of GM and their corresponding eigenvalues for ROI analyses. Partial correlations were calculated between the eigenvalues of each cluster and each of the CDS time points, while accounting for the same models as in the VBM model. The sole purpose of this analysis is to provide further information on whether the other timepoints show similar results that just did not reach the required level of significance.

##### Sensitivity analyses

ROI eigenvalues were extracted using MarsBaR. This approach allowed us to explore the significant clusters from the main analyses while adding potential confounders related to pregnancy and perinatal period. Model 1 additionally controlled for total prenatal distress (sum of EPDS and SCL scores from 14, 24, and 34 GW) at postnatal timepoints (3, 6, and 24 months) and vice versa. Model 2 added potential confounders related to pregnancy and perinatal period. Full models presented in Supplementary Materials.

Additionally, we performed VBM analyses in a which we used all the same covariates as in the main CDS analyses and additionally included the CDS scores from all other timepoints in order to better differentiate the unique contribution of the specific measurement point. Full regression models presented in Supplementary Materials.

## Results

### Voxel-based morphometry results – distress

We observed several significant associations between maternal distress symptoms at various time points during the offspring’s early life and regional GM volumes at 5 years of age. More maternal symptoms at the ages of 3 months and 24 months were associated with smaller volumes, while symptoms at 14 GW and 6 months postpartum were associated with larger volumes. See Figures 1–5 and Table 2.

**Table 2.**
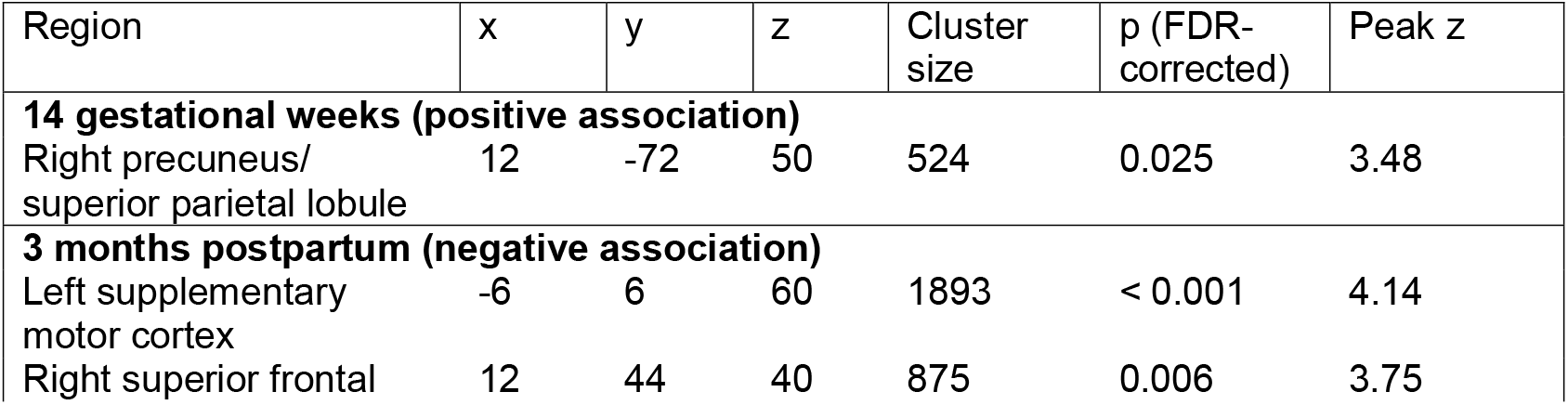
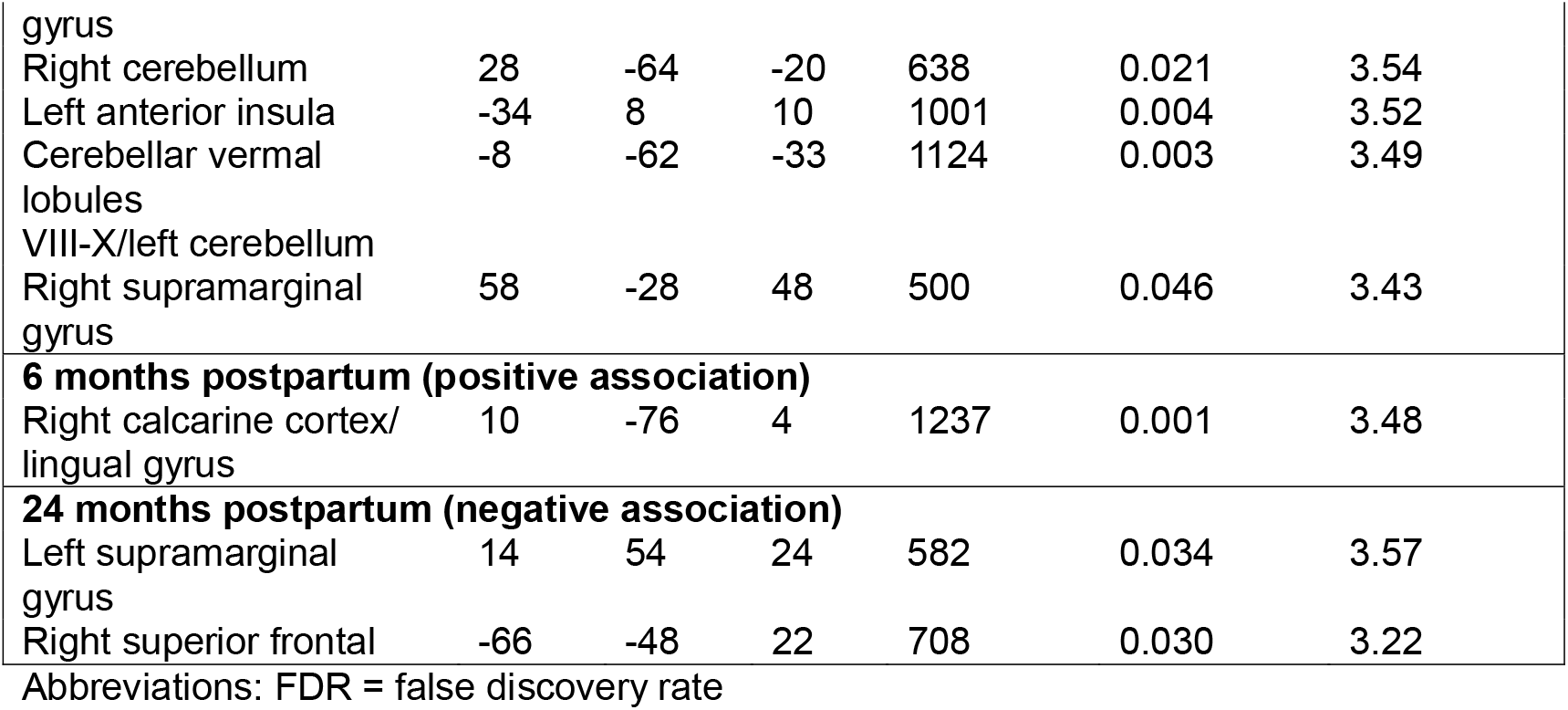
Clusters from the voxel-based morphometry analysis.

**Figure 1.**
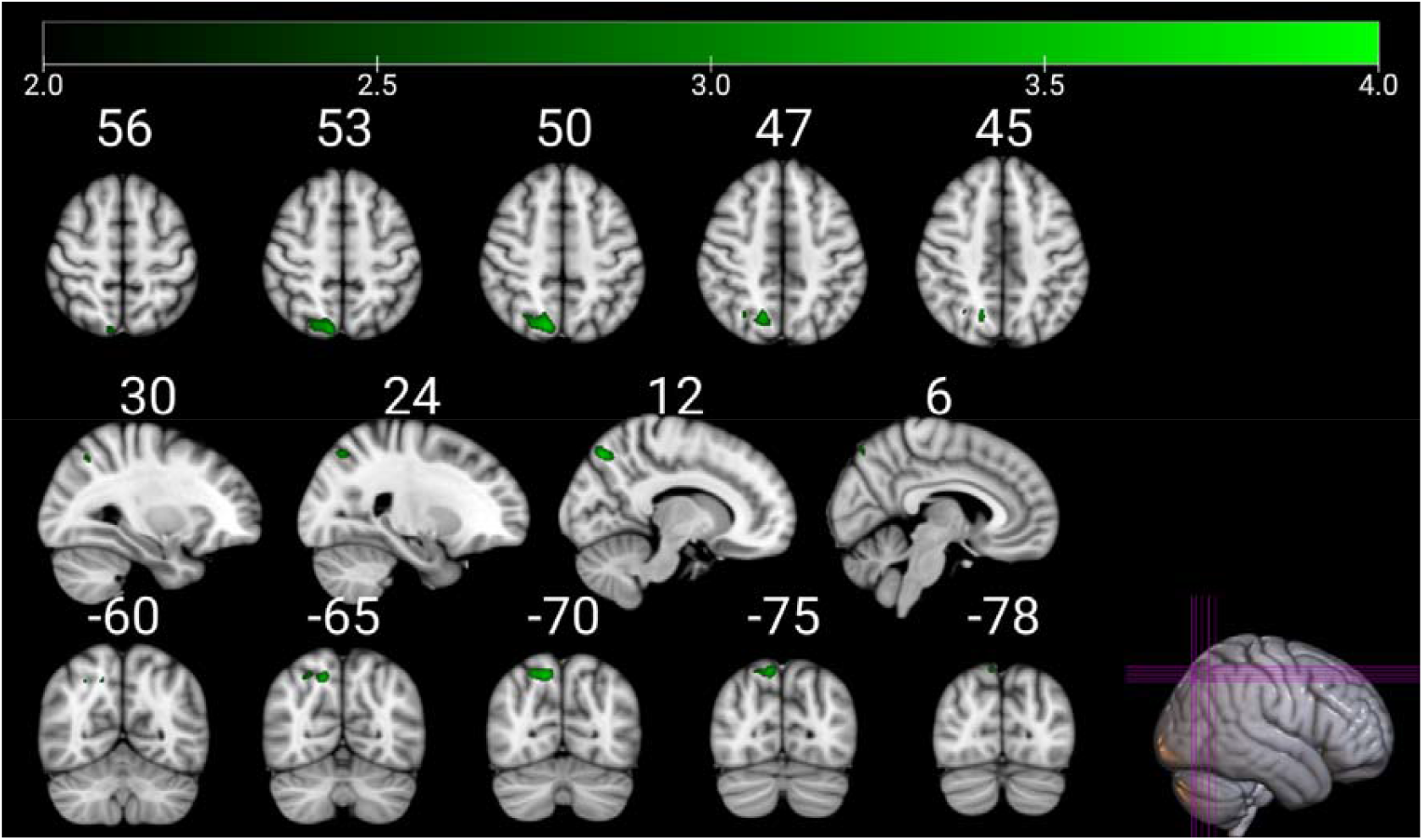
Positive association between maternal distress symptoms at 14 gestational weeks and regional volume in right superior parietal areas, including the parietal lobule and precuneus. Only statistically significant regions at p < 0.005, false discovery rate corrected at cluster level are displayed. Color indicates t-value. Images are in radiological orientation. Coordinates (indicated with white numbers) are in MNI space. The figure was created with MRIcroGL.

**Figure 2.**
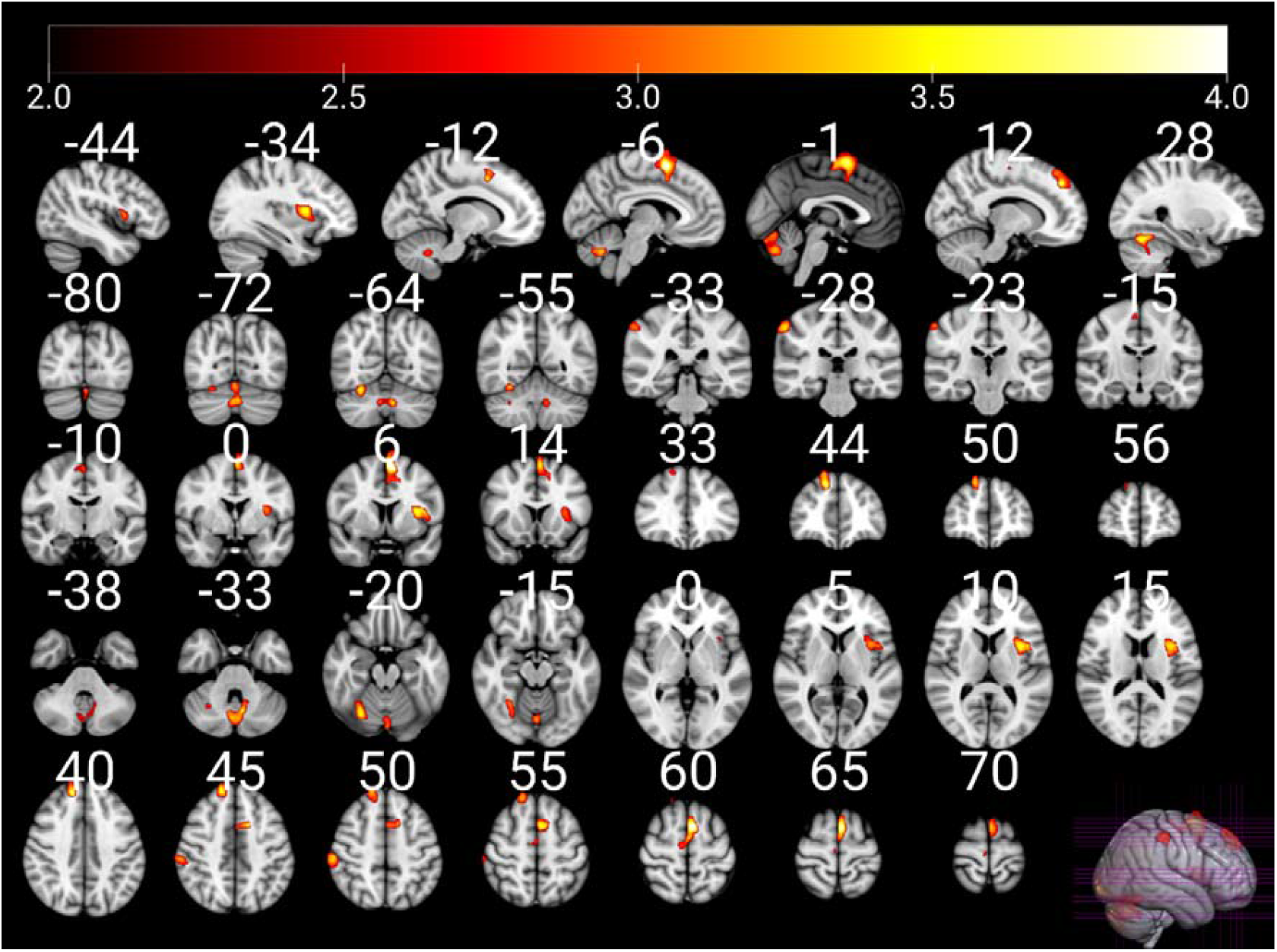
Negative associations between maternal distress symptoms at 3 months postpartum and regional volume in six clusters: right superior frontal gyrus; right cerebellum; right supramarginal gyrus; left cerebellum and cerebellar vermal lobules VIII-X; left supplementary motor cortex; left anterior insula. Only statistically significant regions at p < 0.005, false discovery rate corrected at cluster level are displayed. Color indicates t-value. Images are in radiological orientation. Coordinates (indicated with white numbers) are in MNI space. The figure was created with MRIcroGL.

**Figure 3.**
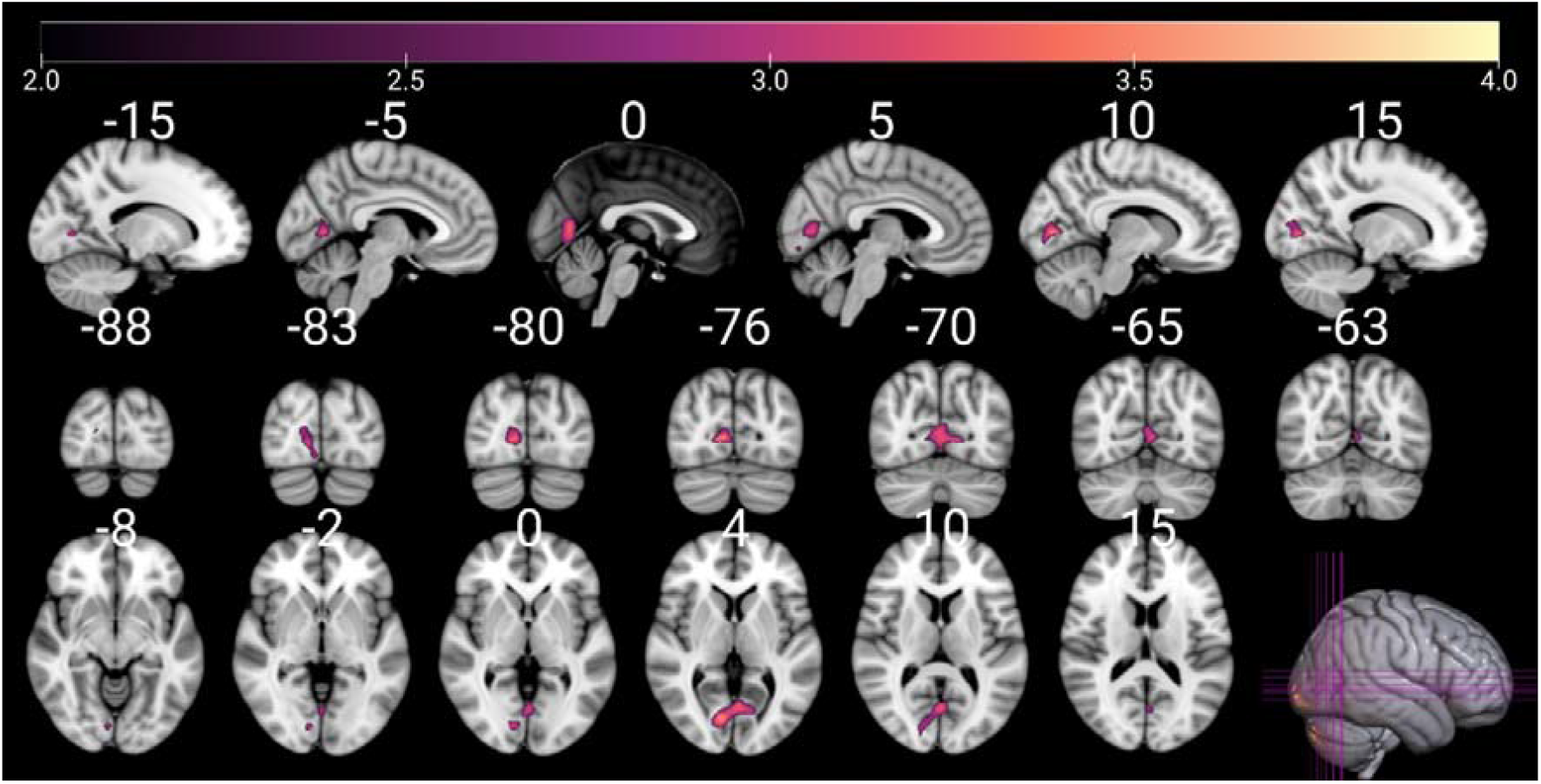
Positive association between maternal distress symptoms at 6 months postpartum and regional volume in the occipital lobe, including the right calcarine cortex and bilateral lingual gyrus. Only statistically significant regions at p < 0.005, false discovery rate corrected at cluster level are displayed. Color indicates t-value. Images are in radiological orientation. Coordinates (indicated with white numbers) are in MNI space. The figure was created with MRIcroGL.

**Figure 4.**
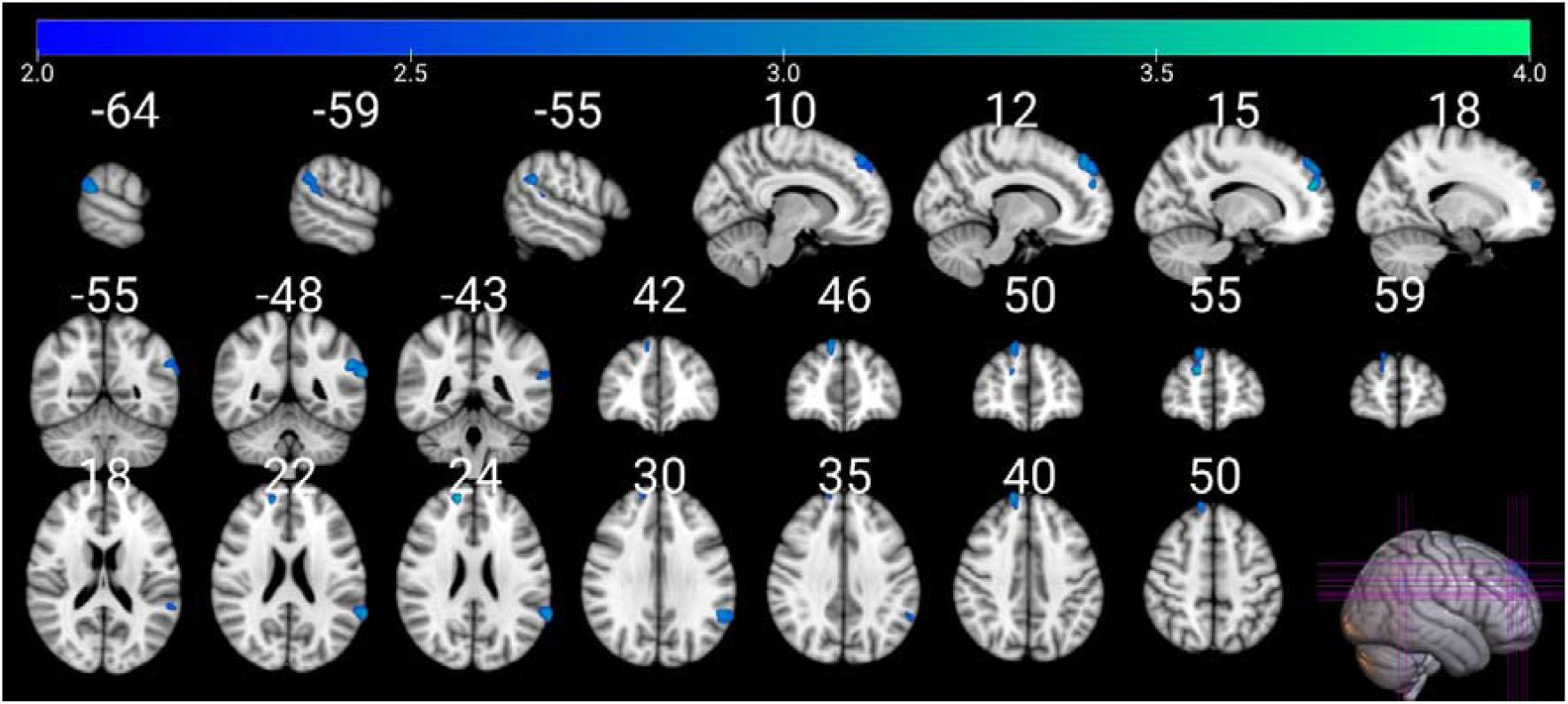
Negative associations between maternal distress symptoms at 24 months postpartum and regional volume in two clusters: the right superior frontal gyrus and left supramarginal gyrus. Only statistically significant regions at p < 0.005, false discovery rate corrected at cluster level are displayed. Color indicates t-value. Images are in radiological orientation. Coordinates (indicated with white numbers) are in MNI space. The figure was created with MRIcroGL.

**Figure 5.**
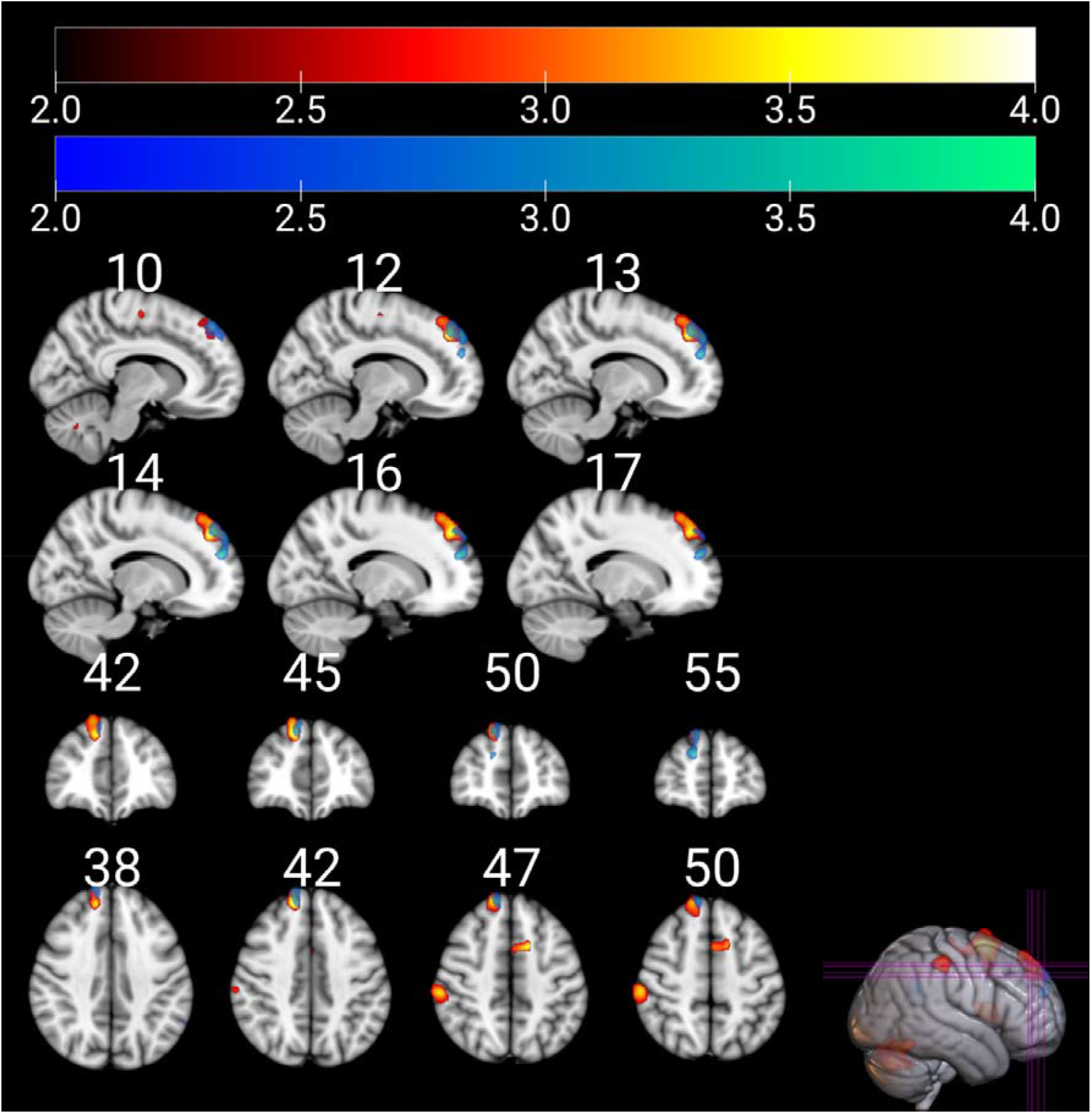
Negative associations between maternal distress symptoms at 3 (hot colormap) and 24 months postpartum (blue–green colormap), highlighting the overlap in superior frontal regions. Only statistically significant regions at p < 0.005, false discovery rate corrected at cluster level are displayed. Color indicates t-value. Images are in radiological orientation. Coordinates (indicated with white numbers) are in MNI space. The figure was created with MRIcroGL.

Same analyses with a CDS based on standardized values resulted in the same number of significant clusters at all same age points, with the same peak coordinates (one exception: cluster (-34, 8, 19) changed to (-33, 8, 12) at 3 months postpartum). Largest difference in cluster size compared to the unstandardized scores was 8%.

### Partial correlation analysis

The partial correlation analysis found significant partial correlations between maternal symptoms and clusters identified in the VBM analysis. Partial correlations are presented in Table 3.

**Table 3.** Partial correlation coefficients (no correction for multiple comparisons) between significant clusters and maternal symptoms at different age points controlling for age at scan, sex, ponderal index, maternal age at term and maternal education.

|  | **CDS**<br>**14 GW** | **CDS**<br>**24 GW** | **CDS**<br>**34 GW** | **CDS**<br>**3 months** | **CDS**<br>**6 months** | **CDS**<br>**24**<br>**months** |
| --- | --- | --- | --- | --- | --- | --- |
| **14 gwks**<br>Right precuneus/<br>superior parietal lobule<br>(12, -72, 50) | .29*** | .17* | .24** | .09 | .08 | .08 |
| **3 months**<br>Left supplementary<br>motor cortex<br>(-6, 6, 60) | -.12 | -.17* | -.06 | -.32*** | -.13 | -.18* |
| **3 months**<br>Cerebellar vermal<br>lobules<br>VIII-X/left cerebellum<br>(-8, -62, -33) | -.16* | -.18* | -.08 | -.29*** | -.19* | -.16* |
| **3 months**<br>Left anterior insula<br>(-34, 8, 10) | -.09 | -.16* | -.05 | -.27*** | -.07 | -.11 |
| **3 months**<br>Right superior frontal<br>gyrus<br>(12, 44, 40) | -.18* | -.11 | -.11 | -.27*** | -.22** | -.19* |
| **3 months**<br>Right cerebellum<br>(28, -64, -20) | -.06 | -.12 | -.02 | -.26*** | -.03 | -.12 |
| **3 months**<br>Right supramarginal<br>gyrus<br>(58, -28, 48) | -.13 | -.16* | -.06 | -.25*** | -.12 | -.18* |
| **6 months**<br>Right calcarine cortex/<br>lingual gyrus<br>(10, -76, 4) | .04 | .05 | .13 | .02 | .28*** | .10 |
| **24 months**<br>Left supramarginal gyrus<br>(-66, -48, 22) | -.08 | -.14 | -.13 | -.18* | -.13 | -.25** |
| **24 months**<br>Right superior frontal<br>gyrus<br>(14, 54, 24) | -.15 | -.07 | -.06 | -.24** | -.16* | -.27*** |

\*\*\* = $p \leq 0.001$ , \*\* = $p \leq 0.01$ , \* = $p \leq 0.05$
CDS = Composite Distress Score, GW = gestational weeks, () = coordinates (x, y, z)

### Voxel-based morphometry results – depressive symptoms

Results for 3 and 6 months of postnatal age were mostly similar to those seen in CDS analyses (Figures S2 and S3). At 34 GW, there were multiple positive associations between maternal depressive symptoms and regional brain volumes: the left cerebellum, right occipital pole, right precentral gyrus, right superior parietal lobule (Figure 6). In contrast to CDS results, using only EPDS score did not yield results at 14 GW or 24 months.

**Figure 6.**
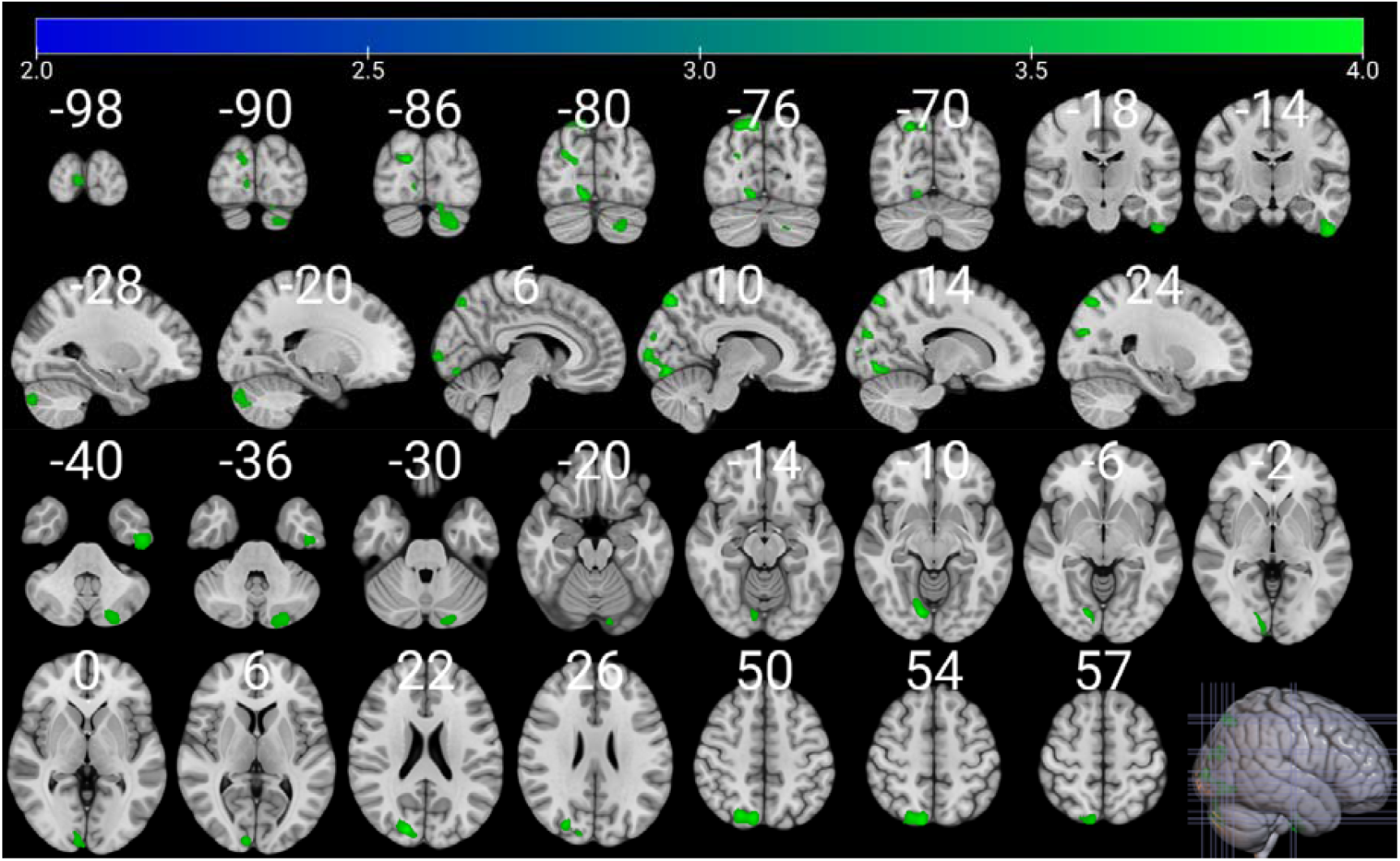
Positive associations between maternal depressive symptoms at 34 gestational weeks and regional volume in four clusters: the left cerebellum, right occipital pole, right precentral gyrus, and right superior parietal lobule. Color indicates t-value. Images are in radiological orientation. Coordinates (indicated with white numbers) are in MNI space. The figure was created with MRIcroGL.

### Sensitivity analyses

In Model 1, all clusters remained significant when adding pre- or postnatal summed CDS to the model. Prenatal CDS only remained in the final model in 2/10 cases (the right cerebellum and the left supplementary motor cortex clusters from 3 months), and it was not statistically significant in either case. In model 2, CDS remained significant for each cluster (p ≤.003 in all cases). All regression models are presented in Tables S3–S24.

In the VBM model in which all CDS timepoints were controlled, multiple significant positive results were observed between brain volume and CDS at 34 GW and 6 months postpartum, and multiple negative results between brain volume and 3 months postpartum. Results from 3 and 6 months postpartum partially overlap with the main VBM models. Results are presented in Table S25 and Figure S4.

## Discussion

This study investigated the association between pre- and postnatal maternal distress and child brain development at 5 years of age in the FinnBrain Birth Cohort study. We found that prenatal and particularly early postnatal symptoms of depression and anxiety were associated with offspring brain morphology. The most prominent findings were found at 3 months postnatal age. The associations with maternal distress were negative at 3 and 24 months, and positive at 14 GW and 6 months. At 34 GW, significant results were only found for depressive symptoms, not overall distress exposure. Contrary to our hypotheses, no significant results were found in the hippocampus or amygdala.

Prenatally, only 14 GW showed significant associations with distress, located in the right superior parietal regions, including the precuneus. The superior parietal region is involved in various cognitive functions, including visuospatial processing and working memory [59], while the precuneus is involved in memory retrieval and attentional processing [60]. The right superior parietal region [61,62] and precuneus [61] surface area and CT have been associated with prenatal distress exposure before. Overall, the regions associated with early gestation exposure are important for cognitive functions [63], and our earlier study in the same population found an association between non-verbal cognitive ability and cortical volume and surface in a cluster involving the precuneus [64]. Finally, the scarcity of prenatal findings in our study was surprising, considering the previous literature [2].

Postnatal exposure to maternal distress was associated with widespread structural changes in the offspring’s brain. The superior frontal and supramarginal gyrus [65], as well as the cerebellum [66– 68] are regions involved in cognitive processes, including semantic processing and social cognition [69]. The cerebellar findings in this study involved both motor and non-motor regions [70]. Together with the prenatal results, these findings indicate a possible connection between prenatal and early life distress exposure and cognitive development. This connection has been studied following the 1998 ice storm in Québec, Canada [71], wherein prenatal distress exposure explained over 10% of the variance in general cognitive ability and language skills in toddlerhood. Motor control is another aspect that connects to many of the regions in our study: the cerebellum, left supplementary motor cortex, and left anterior insula [72]. Motor development has also been studied in the context of the Québec ice storm and there is some evidence of prenatal distress exposure affecting motor development in children [73]. Our results indicate a potential connection between prenatal and early life distress exposure with cognitive and motor development, which will be important questions for future studies.

The hippocampus and amygdala have been associated with prenatal distress in multiple prior studies. Postnatal distress literature is more limited [17,21,27]. We found no significant clusters in either one associated with distress or depressive symptoms at any timepoint. The previous results have been inconsistent, and might be affected by timing of exposure, or the research questions (e.g., association only when looking at different genetic profiles [32] or sexes [22] separately). Furthermore, hippocampus and particularly the amygdala are small structures that are challenging to segment accurately [74,75], and methodological variation could also explain some of the inconsistent results.

In this study, we tested the neural correlates of distress and depressive symptoms separately. For postnatal analyses, the results were quite similar: multiple significant negative associations at 3 months and one cluster in the calcarine and lingual regions at 6 months. The were no clusters in new regions and the decrease in significant results was expected as the variance decreased – particularly considering that the symptoms were mostly in the normal range in our sample. For the prenatal analyses, there was a meaningful difference between distress and depressive symptoms. In the third trimester, depressive symptoms were associated with brain volumes in multiple clusters, while distress was not. The results were similar in the sense that postnatal age points produced more results than prenatal ones, and the seemingly large difference at 34 GW could be partially related to thresholding.

Three months of postnatal age was the time that most significant associations were observed. This was the case for both distress and depressive symptoms, the findings persisted (although some regions changed) when controlling for all other age points, and in the supplementary stepwise regression models adding prenatal distress in the model for postnatal results and vice versa rarely remained in the model and was never statistically significant. However, it is important to note that the repeated distress measurements are highly intercorrelated, and this modelling approach does not capture information about the longitudinal exposure to maternal distress. A trajectory-based approach was considered but based on our prior research [78] we know that trajectories showing mostly prenatal or mostly postnatal symptoms are rare in the FinnBrain Birth Cohort. The early postnatal life is a potential sensitive period for neural development, as it is characterized by neural processes like myelination and synaptogenesis [76] as well as quick increase in cerebral GM volume [77]. Our results highlight the need to study the early postnatal period further, rather than just pregnancy.

This study has some limitations. Questionnaire data was collected longitudinally, with slightly increasing amount of missing data towards later measurements. This study relied on self-report questionnaires to assess maternal anxiety and depressive symptoms, which can be subject to measurement errors. Despite the potential issues, self-report remains the most common and widely accepted method for collecting data on subjective experiences, such as depressive and anxiety symptoms. The questionnaire data was analyzed at multiple cross-sectional timepoints and not as longitudinal trajectories as discussed above. Finally, the homogeneity of our sample, consisting of FinnBrain participants mainly of Finnish origin from a specific region (i.e., Southwestern Finland), with a generally higher level of education, limits the generalizability of our findings to other populations and contexts.

## Conclusion

This study investigated the associations between maternal distress during the pre- and postnatal periods and the brain development of offspring at the age of 5 years. A particular strength of our work is the collection of distress measurements from multiple timepoints both pre- and postnatally. Our findings revealed connections between exposure to maternal distress at various developmental stages and structural alterations in widespread brain regions, particularly those associated with cognitive and motor functions. In contrast to prior literature, this study highlights the importance of the early postnatal period as a sensitive window for structural brain development. Importantly, structural associations with early life distress do not automatically indicate maladaptive development. These results form a basis for hypothesis creation for developmental pathways from early distress exposure to cognitive or behavioral outcomes. In conclusion, future studies should always consider the timing of the exposure, including the postnatal period, to the extent that they have available data.

## Acknowledgements

We thank the FinnBrain Birth Cohort study participants, the staff, and the assisting personnel. We thank our research nurse Susanne Sinisalo, neuroradiologist Riitta Parkkola, paediatric neurologist Tuire Lähdesmäki, and physicist Jani Saunavaara for their role in the 5-year MRI data collection.

## Supplementary material

For supplementary material accompanying this paper, visit cambridge.org/EPA.

