## Supplementary Materials for "Exposure to Maternal Pre- and Postnatal Psychological Distress: Associations with Brain Structure in 5-year-old Children"

### Methods and Materials

This study follows the Strengthening the Reporting of Observational Studies in Epidemiology (STROBE) guidelines (1), see Table S1 for the attached checklist.

Table S1. STROBE checklist

|  | Item No | Recommendation | Location |
| --- | --- | --- | --- |
| **Title and abstract** | 1 | (*a*) Indicate the study’s design with a commonly used term in the title or the abstract | Abstract |
|  |  | (*b*) Provide in the abstract an informative and balanced summary of what was done and what was found | Abstract |
| Introduction | | | |
| Background/rationale | 2 | Explain the scientific background and rationale for the investigation being reported | Introduction |
| Objectives | 3 | State specific objectives, including any prespecified hypotheses | The last paragraph of “Introduction” |
| Methods | | | |
| Study design | 4 | Present key elements of study design early in the paper | Procedures |
| Setting | 5 | Describe the setting, locations, and relevant dates, including periods of recruitment, exposure, follow-up, and data collection | Participants and Supplementary materials |
| Participants | 6 | (*a*) Give the eligibility criteria, and the sources and methods of selection of participants | Participants and Supplementary materials |
| Variables | 7 | Clearly define all outcomes, exposures, predictors, potential confounders, and effect modifiers. Give diagnostic criteria, if applicable | Measures and Table 1 footnotes |
| Data sources/ measurement | 8 | For each variable of interest, give sources of data and details of methods of assessment (measurement). Describe comparability of assessment methods if there is more than one group | Measures and Image acquisition |
| Bias | 9 | Describe any efforts to address potential sources of bias | Supplementary materials |
| Study size | 10 | Explain how the study size was arrived at | Participants and Supplementary materials |
| Quantitative variables | 11 | Explain how quantitative variables were handled in the analyses. If applicable, describe which groupings were chosen and why | Data preprocessing and statistical analyses |
| Statistical methods | 12 | (*a*) Describe all statistical methods, including those used to control for confounding | Data preprocessing and statistical analyses |
|  |  | (*b*) Describe any methods used to examine subgroups and interactions | N/A |
|  |  | (*c*) Explain how missing data were addressed | Measures |
|  |  | (*d*) If applicable, describe analytical methods taking account of sampling strategy | N/A |
|  |  | (*e*) Describe any sensitivity analyses | Statistics |
| Results | | | |
| Participants | 13 | (a) Report numbers of individuals at each stage of study—eg numbers potentially eligible, examined for eligibility, confirmed eligible, included in the study, completing follow-up, and analysed | Supplementary materials |
|  |  | (b) Give reasons for non-participation at each stage | N/A |
|  |  | (c) Consider use of a flow diagram | N/A |
| Descriptive data | 14 | (a) Give characteristics of study participants (eg demographic, clinical, social) and information on exposures and potential confounders | Table 1 |
|  |  | (b) Indicate number of participants with missing data for each variable of interest | Measures and Table 1 |
| Outcome data | 15 | Report numbers of outcome events or summary measures | Table 1 and Supplementary materials |
| Main results | 16 | (*a*) Give unadjusted estimates and, if applicable, confounder-adjusted estimates and their precision (eg, 95% confidence interval). Make clear which confounders were adjusted for and why they were included | No unadjusted estimates. Main adjusted results under “Results”, confidence intervals in regression models included in Supplementary materials. |
|  |  | (*b*) Report category boundaries when continuous variables were categorized | N/A |
|  |  | (*c*) If relevant, consider translating estimates of relative risk into absolute risk for a meaningful time period | N/A |
| Other analyses | 17 | Report other analyses done—eg analyses of subgroups and interactions, and sensitivity analyses | Results and Supplementary materials |
| Discussion | | | |
| Key results | 18 | Summarise key results with reference to study objectives | First paragraph of Discussion |
| Limitations | 19 | Discuss limitations of the study, taking into account sources of potential bias or imprecision. Discuss both direction and magnitude of any potential bias | Last paragraph of Discussion |
| Interpretation | 20 | Give a cautious overall interpretation of results considering objectives, limitations, multiplicity of analyses, results from similar studies, and other relevant evidence | Discussion |
| Generalisability | 21 | Discuss the generalisability (external validity) of the study results | Last paragraph of Discussion |
| Other information | | | |
| Funding | 22 | Give the source of funding and the role of the funders for the present study and, if applicable, for the original study on which the present article is based | Acknowledgements |

#### Participants

The participants are a part of the prospective FinnBrain Birth Cohort Study ([www.finnbrain.fi](http://www.finnbrain.fi)), which examines the influence of genetic and environmental factors on child development and later health outcomes (2). Pregnant women (n = 3808) attending their first trimester ultrasound at gestational week (GW) 12, their spouses (n = 2623), and children (n = 3837; including 29 twin pairs) were recruited in Southwest Finland between December 2011 and April 2015. An ultrasound-verified pregnancy and sufficient knowledge of Finnish or Swedish languages were required for participation. The cohort study includes several follow-up studies.

For the neuroimaging visit, we primarily recruited participants that had attended the neuropsychological visit. The participants recruited for both visits were focus cohort families (2) and families who had actively participated in previous FinnBrain study visits. For the neuroimaging visits, 541 families were contacted and 478 (88.4%) of them were reached. In total, 203 (37.5%) participants attended imaging visits (113 boys (55.7%), mean age 5.40 (SD 0.13), range 5.08–5.79 years). For this study, we included only participants with an adequate quality T1 image, assessed by the author E.P.P. in our prior work (3), resulting in the final sample size of 173.

We originally aimed to scan all subjects between the ages 5 years 3 months and 5 years 5 months; however, there was a pause in visits due to the start of the COVID-19 pandemic, and subsequently many of the participants were older than planned when they were scanned (152/203 (74.9%) of the participants attended the visit within the intended age range). The exclusion criteria for the neuroimaging study were: 1) born before GW 35 (before GW 32 for those with exposure to maternal prenatal synthetic glucocorticoid treatment), 2) developmental anomaly or abnormalities in senses or communication (e.g., blindness, deafness, congenital heart disease), 3) known long-term medical diagnosis (e.g., epilepsy, autism), 4) ongoing medical examinations or clinical follow up in a hospital (meaning there has been a referral from primary care setting to special health care), 5) child use of continuous, daily medication (including per oral medications, topical creams and inhalants; One exception to this was desmopressin medication, which was allowed), 6) history of head trauma (defined as concussion necessitating clinical follow up in a health care setting or worse), 7) metallic (golden) ear tubes (to assure good-quality scans), and routine magnetic resonance imaging (MRI) contraindications.

#### Bias assessment

Mothers of the children who did not participate in the neuropsychological visits (out of the 1288 contacted families) had a lower education level (χ2(2) = 30.94, p < 0.001), a lower monthly income (χ2(3) = 11.65, p = 0.009) and were younger (t (1286) = -4.130, p < 0.001) compared to the mothers in the families that participated in the neuropsychological visits.

Mothers of the children who participated in the neuropsychological visits but not in the neuroimaging visits were older (t (369) = 1.97, p = 0.047) but did not differ in education level or monthly income compared to the mothers in the families that participated in the MRI visit.

#### Measures

**Depressive symptoms**: To assess maternal depressive symptoms, the Finnish version of the Edinburgh Postnatal Depression Scale (EPDS)(4) was used. EPDS is a 10-item self-report screening tool that is widely used to identify depressive symptoms in women both during and after pregnancy. The participant is asked to report their symptoms from the past seven days. Each item is rated on a 4-point Likert scale (0 = never or not at all, 3 = very often or most of the time), with total scores ranging from 0 to 30. The clinical cut-off threshold for potential depression is ≥ 10 points.

**Anxiety symptoms**: Maternal anxiety symptoms were evaluated using the Finnish version of the Symptom Checklist-90-Revised (SCL)(5). SCL is a self-report questionnaire comprising of 90 questions, designed to assess a wide range of psychiatric and somatic symptoms. However, for the purposes of this study, only the anxiety subscale was utilized. The 10-item scale assesses signs of nervousness, tension, and feelings of panic. Items are scored on a 5-point scale assessing the level of distress a given symptom has caused in the past month (0 = not at all, 4 = extremely), with total scores ranging from 0 to 40 and higher scores indicating more symptoms.

The EPDS and SCL scores were not standardized while calculating the CDS, as that would make the effect sizes not comparable with other studies (Pearson correlations between non-standardized and standardized CDS = 1.000 at each timepoint; Spearman correlation > 0.998 at each timepoint).

A correlation matrix of EPDS and SCL scores from all age points is presented in Table S2. Missing data is visualized in Figure S1.

Table S2. Spearman’s rho correlations between maternal depressive and anxiety symptoms at different age points

|  | **EPDS 14 GW** | **EPDS 24 GW** | **EPDS 34 GW** | **EPDS 3 mo** | **EPDS 6 mo** | **EPDS 24 mo** | **SCL 14 GW** | **SCL 24 GW** | **SCLs 34 GW** | **SCL 3 mo** | **SCL 6 mo** |
| --- | --- | --- | --- | --- | --- | --- | --- | --- | --- | --- | --- |
| **EPDS 14 GW** | -- |  |  |  |  |  |  |  |  |  |  |
| **EPDS 24 GW** | .557^***^ | -- |  |  |  |  |  |  |  |  |  |
| **EPDS 34 GW** | .541^***^ | .644^***^ | -- |  |  |  |  |  |  |  |  |
| **EPDS 3 mo** | .456^***^ | .507^***^ | .428^***^ | -- |  |  |  |  |  |  |  |
| **EPDS 6 mo** | .449^***^ | .477^***^ | .393^***^ | .654^***^ | -- |  |  |  |  |  |  |
| **EPDS 2y** | .295^**^ | .327^***^ | .374^***^ | .531^***^ | .538^***^ | -- |  |  |  |  |  |
| **SCL 14 GW** | .698^***^ | .448^***^ | .386^***^ | .371^***^ | .289^***^ | .197^*^ | -- |  |  |  |  |
| **SCL 24 GW** | .579^***^ | .680^***^ | .516^***^ | .491^***^ | .426^***^ | .262^**^ | .674^***^ | -- |  |  |  |
| **SCL 34 GW** | .502^***^ | .497^***^ | .601^***^ | .406^***^ | .372^***^ | .243^**^ | .649^***^ | .733^***^ | -- |  |  |
| **SCL 3 mo** | .433^***^ | .452^***^ | .381^***^ | .594^***^ | .419^***^ | .305^***^ | .536^***^ | .554^***^ | .581^***^ | -- |  |
| **SCL 6 mo** | .435^***^ | .491^***^ | .449^***^ | .591^***^ | .717^***^ | .404^***^ | .457^***^ | .523^***^ | .579^***^ | .661^***^ | -- |
| **SCL 24 mo** | .266^**^ | .294^***^ | .298^***^ | .435^***^ | .443^***^ | .592^***^ | .376^***^ | .327^***^ | .357^***^ | .460^***^ | .457^***^ |

*** = p < 0.001, ** = p < 0.01, * = p < 0.05  
EPDS = Edinburgh Postnatal Depression Scale, SCL = Symptom Checklist, gwks = gestational weeks, mo = months. Spearman correlations calculated without data imputation. Number of missing data points: 14 GW n = 9, 24 GW n = 8, 34 GW n = 8; 3 months n = 17, 6 months n = 32, 24 months n = 47

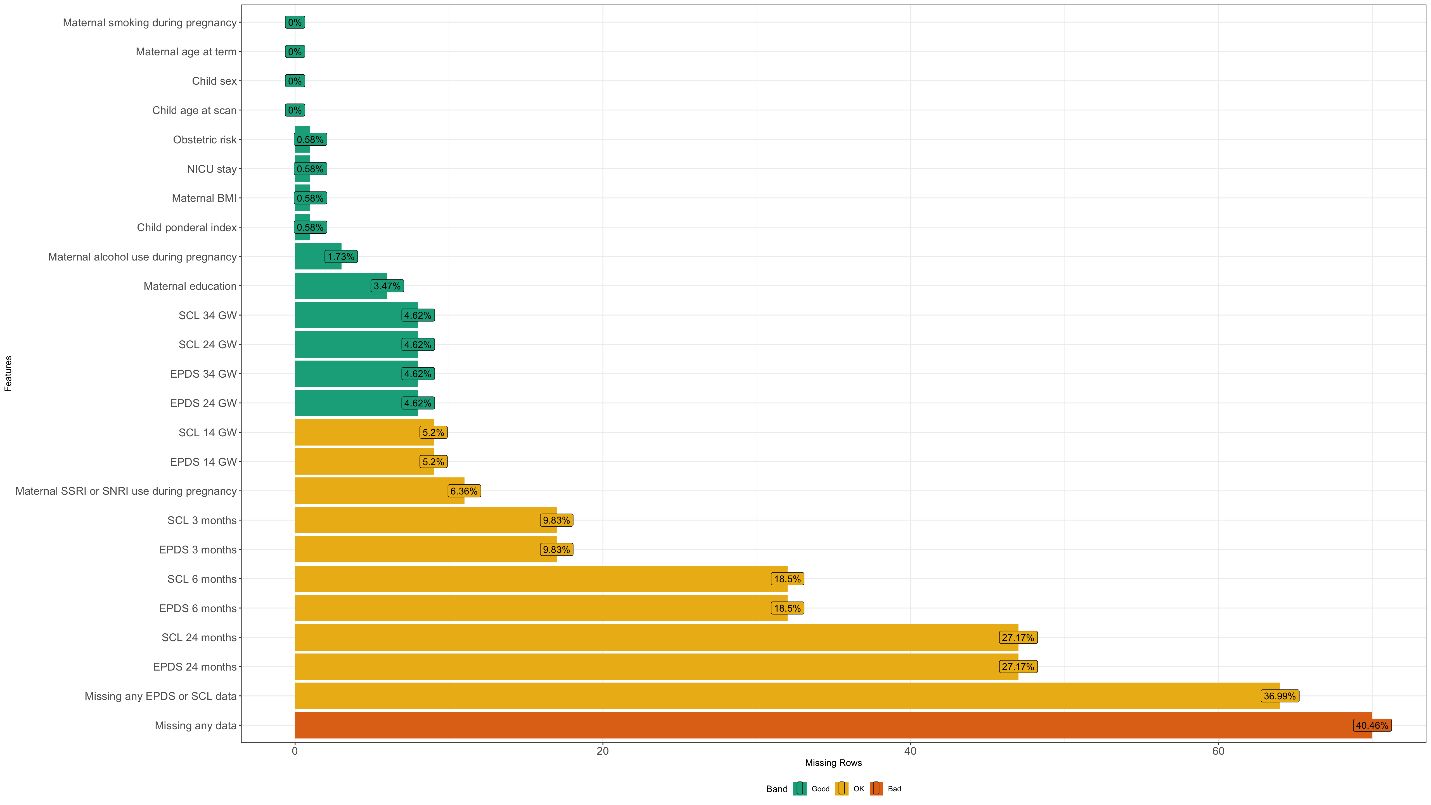

Figure S1. Visualization of all the missing data in the final analysis sample (173 participants).

#### Neuroimaging visits

MRI scanning was conducted at Turku University Hospital by research staff. Prior to the visits, families were contacted and the purpose of the study, along with the scanning procedure, were explained to the parents. Parents were then requested to share this information with their child and inquire whether they would like to participate. Follow-up phone calls were made by a research nurse to confirm the willingness to participate. Written informed consent was obtained from both parents prior to any experimental procedures. In preparation for the MRI imaging visits, a member of the research team met with each participating family in person. The families were provided with practice materials to help the children become familiar with the imaging procedure and to facilitate practice sessions for scanning at home. On the day of the MRI, a simulated scan was conducted in the laboratory using a homemade non-commercial wooden coil to further familiarize the child with the scanning process. Additionally, as is customary in MRI studies involving children, a light meal was provided before the actual experimental scans took place (6).

Participants underwent scanning either during natural sleep or awake while watching a movie/TV show of their preference. Earplugs and headphones were provided for hearing protection, and foam padding was applied to reduce head movement and enhance comfort during scanning. Throughout the scanning process, a member of the research team and the child’s parent(s) stayed in the scanner room. Participants were given a signal ball that they could use to pause or stop the scan at any point if needed. More detailed information regarding the protocol and study visits is presented in previous publications (3,7).

#### Statistics

**ROI-based sensitivity analyses:**

ROI eigenvalues were extracted using MarsBaR. This approach allowed us to explore the significant clusters from the main analyses while adding covariates. To explore the specificity of the structural brain differences for pre- or postnatal distress, we performed regression models, in which we additionally controlled for total prenatal distress (sum of EPDS and SCL scores from 14, 24, and 34 GW) at postnatal timepoints (3, 6, and 24 months) and vice versa. Stepwise regression model using the Akaike information criterion (forward and backward) was used to optimize the variables left in the final regression model.

*Model 1a: Prenatal ROI eigenvalues ~ Age at scan + Sex + Ponderal Index + Maternal age at term + Education + Composite Distress Score from the ROI timepoint + postnatal Composite Distress Score*

*Model 1b: Postnatal ROI eigenvalues ~ Age at scan + Sex + Ponderal Index + Maternal age at term + Education + Composite Distress Score from the ROI timepoint + prenatal Composite Distress Score*

Additionally, we used the extracted eigenvalues to explore if the results remain after potential confounders related to pregnancy and perinatal period:

*Model 2: ROI eigenvalues ~ CDS from the time point in which the ROI was significant + Age at scan + Sex + Ponderal Index + Maternal age at term + Maternal education + Prenatal alcohol exposure + Prenatal smoking exposure + Prenatal selective serotonin reuptake inhibitor or serotonin and norepinephrine reuptake inhibitor exposure + Obstetric risk + Maternal pre-pregnancy body mass index + Neonatal intensive care unit admission*

**Voxel-based morphometry sensitivity analyses:**

We performed voxel-based morphometry (VBM) analyses in a which we used all the same covariates as in the main composite distress score (CDS) analyses and additionally included the CDS scores from all other timepoints. For example, at gestational week (GW) 14, we included the following CDS scores: 1) GW 14, 2) sum of GW 24 and GW 34, and 3) sum of all postnatal time points. This was done to limit the effects having multiple highly correlated variables. Variance inflation factors (VIF) were calculated for each model, and were < 2.7 for all variables in all models (data not shown). Full models below:

Model for GW 14:

*VBM contrast ~ Age at scan + Sex + Ponderal Index + Maternal age at term + Maternal education + 14 GW CDS + sum of 24 and 34 GW CDS + postnatal CDS*

Model for GW 24:

*VBM contrast ~ Age at scan + Sex + Ponderal Index + Maternal age at term + Maternal education + 24 GW CDS + sum of 14 and 34 GW CDS + postnatal CDS*

Model for GW 34:

*VBM contrast ~ Age at scan + Sex + Ponderal Index + Maternal age at term + Maternal education + 34 GW CDS + sum of 14 and 24 GW CDS + postnatal CDS*

Model for postnatal month 3:

*VBM contrast ~ Age at scan + Sex + Ponderal Index + Maternal age at term + Maternal education + 3 months CDS + sum of 6 and 24 months CDS + prenatal CDS*

Model for postnatal month 6:

*VBM contrast ~ Age at scan + Sex + Ponderal Index + Maternal age at term + Maternal education + 6 months CDS + sum of 3 and 24 months CDS + prenatal CDS*

Model for postnatal month 24:

*VBM contrast ~ Age at scan + Sex + Ponderal Index + Maternal age at term + Maternal education + 24 months CDS + sum of 3 and 6 months CDS + prenatal CDS*

### Results

#### Voxel-based morphometry results – depressive symptoms

Compared to CDS results, using only EPDS score did not yield results at 14 GW or 2 years. Results for 3 and 6 months of postnatal age were mostly similar to those seen in CDS (Figures S2 and S3). At 34 GW, there were multiple positive associations between maternal depressive symptoms and regional brain volumes: the left cerebellum, right occipital pole, right precentral gyrus, right superior parietal lobule (Figure 6 in the main manuscript).

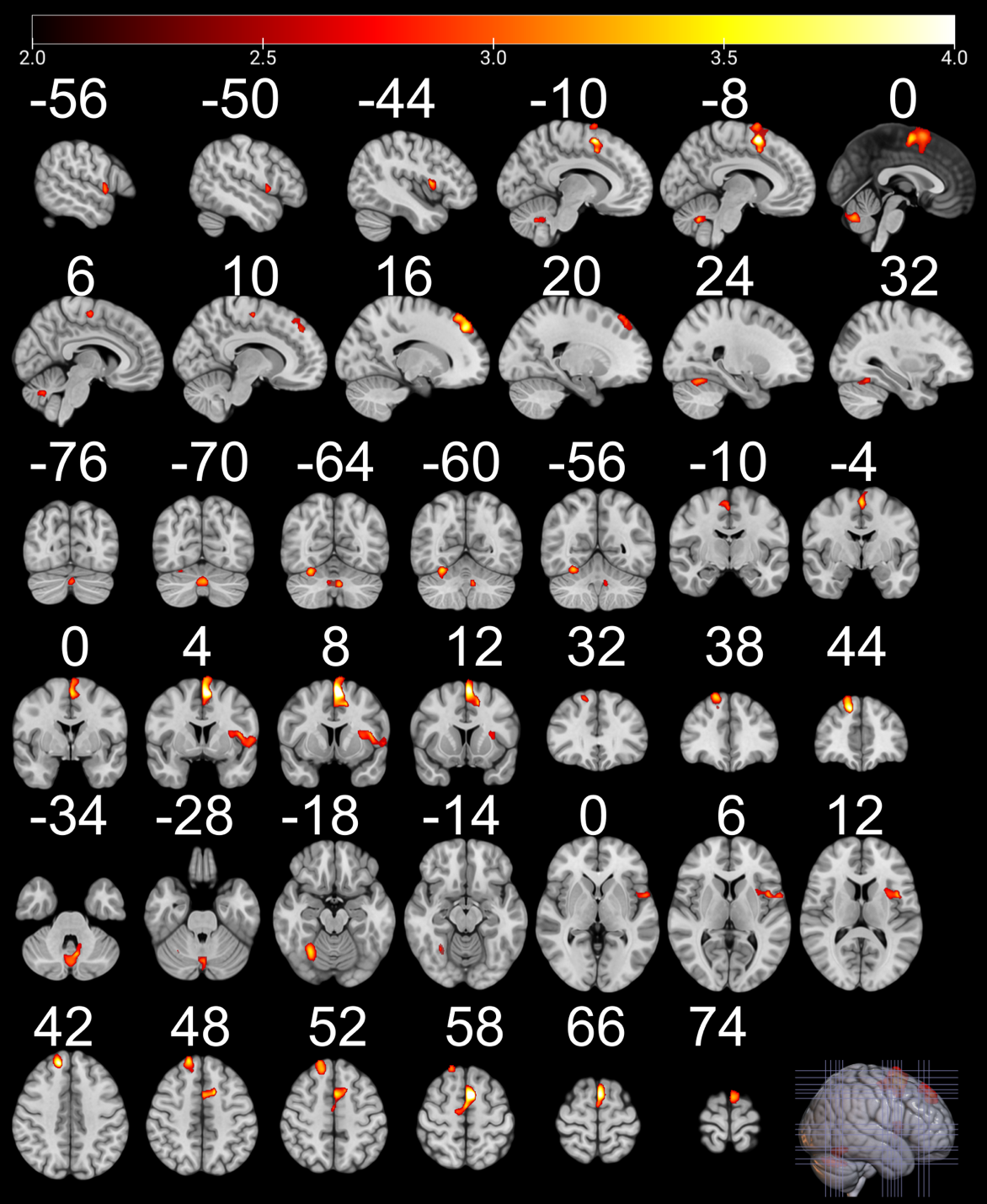

Figure S2. Negative associations between maternal depressive symptoms at 3 months postnatal and regional volume in multiple brain regions: bilateral supplementary motor cortex, left precentral gyrus, left insula, left superior temporal pole, right superior frontal gyrus, right cerebellum, vermis, and cerebellar vermal lobules VI–VII. Only statistically significant regions at p < 0.005, false discovery rate corrected at cluster level are displayed. Color indicates t-value. Images are in radiological orientation. Coordinates (indicated with white numbers) are in MNI space. The figure was created with MRIcroGL.

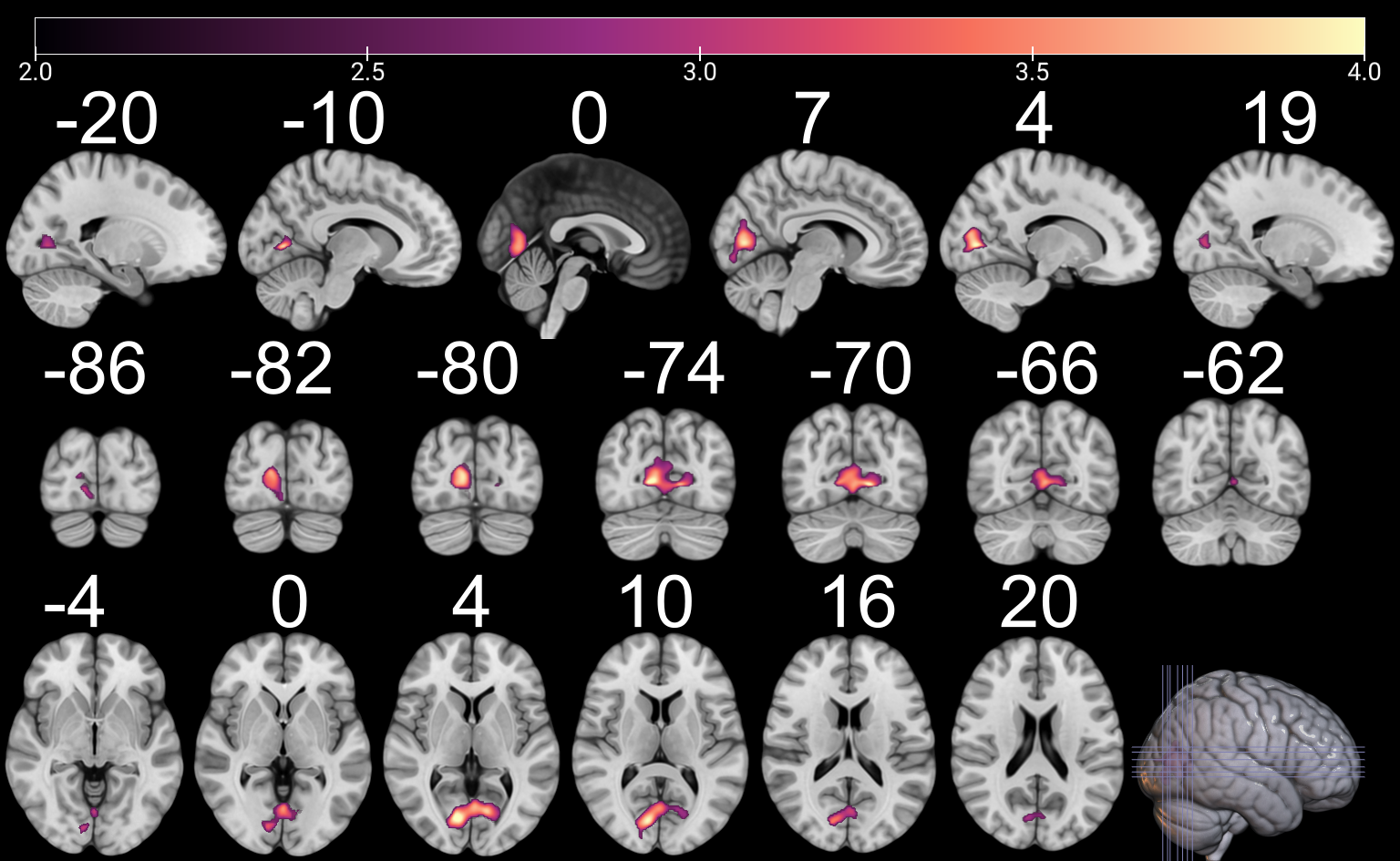
Figure S3. Positive associations between maternal depressive symptoms at 6 months postnatal and regional volume in the occipital lobe, including the bilateral calcarine cortex, bilateral lingual gyrus, left inferior temporal gyrus, and left fusiform gyrus. Only statistically significant regions at p < 0.005, false discovery rate corrected at cluster level are displayed. Color indicates t-value. Images are in radiological orientation. Coordinates (indicated with white numbers) are in MNI space. The figure was created with MRIcroGL.

#### Sensitivity analyses

##### ROI-based analyses

For model 1a, the cluster associated with CDS at 14 GW was significant, F (5,167) = 7.91, p < .001, R2 = 0.19, adjusted R2 = 0.17. The individual significant predictors included: sex (females compared to males) (β = -.20, p = .006), ponderal index (β = .16, p = .02), maternal age at term (β = .18, p = .01), and CDS at 14 GW (β = .27, p < .001). For model 1b, all clusters associated with postnatal CDS regression models were significant. *See Table S3 for a summary of the models*. The significant individual predictors included: sex and the corresponding CDS timepoint for all nine clusters, age at scan for a cluster at 3 months (region of interest [ROI] 1), maternal age at term for a cluster at 24 months (ROI 2), and ponderal index for a cluster at 24 months (ROI 1). *See Tables S4–S13 for the full models*.

In Model 2, all clusters within the models were significant (*see Table S14 for a summary of all models*). Significant individual predictors included: sex (females compared to males) for all 10 clusters, ponderal index for the cluster at 14 GW and clusters at 24 months (ROI 1), age at scan for a cluster at 3 months (ROI 1). Maternal age at term was a significant predictor for clusters at 14 GW and at 24 months (ROI 2), maternal university education for a cluster at 3 months (ROI 4), maternal smoking for clusters at 3 months (ROI 5) and 24 months (ROI 1). NICU admittance was a significant predictor for a cluster at 3 months (ROI 3). Obstetric risk was a significant predictor for a cluster at 3 months (ROI 6). *See Tables S15–S24 for the full models.*

Table S3. Summary of models 1a and b fit statistics for region of interest eigenvalues associated with postnatal CDS additionally controlling for the either prenatal or postnatal CDS.

|  | **F (df1, df2)** | ***p*** | **R^2^** | **Adjusted R^2^** |
| --- | --- | --- | --- | --- |
| **14 gestational weeks** Right precuneus/ superior parietal lobule **(12, -72, 50)** | 7.91*** (5, 167) | <.001 | .19 | .17 |
| **3 months (ROI 1)** Left supplementary motor cortex  **(-6, 6, 60)** | 11.24*** (5, 167) | <.001 | .25 | .23 |
| **3 months (ROI 2)** Cerebellar vermal lobules VIII-X/left cerebellum  **(-8, -62, -33)** | 12.04*** (2, 170) | <.001 | .12 | .11 |
| **3 months (ROI 3)** Left anterior insula  **(-34, 8, 10)** | 20.67*** (2, 170) | <.001 | .20 | .19 |
| **3 months (ROI 4)** Right superior frontal gyrus  **(12, 44, 40)** | 8.86*** (4, 168) | <.001 | .17 | .15 |
| **3 months (ROI 5)** Right cerebellum **(28, -64, -20)** | 22.82*** (3, 169) | <.001 | .29 | .28 |
| **3 months (ROI 6)** Right supramarginal gyrus  **(58, -28, 48)** | 10.68*** (2, 170) | <.001 | .11 | .10 |
| **6 months**  Right calcarine cortex/ lingual gyrus **(10, -76, 4)** | 8.20*** (3, 169) | <.001 | .14 | .11 |
| **24 months (ROI 1)** Left supramarginal gyrus  **(-66, -48, 22)** | 9.32*** (3, 169) | <.001 | .14 | .13. |
| **24 months (ROI 2)**  Right superior frontal gyrus **(14, 54, 24)** | 8.92*** (5, 167) | <.001 | .21 | .19. |

*** = p ≤ 0.001, ** = p ≤ 0.01, * = p ≤ 0.05

CDS = Composite Distress Score, () = coordinates (x, y, z)

Table S4. Regression results using model 1a and 14 gestational weeks ROI (12, -72, 50) as the criterion

| **Predictor** | **b** | **b**  **95% CI**  **[LL, UL]** | **β** | **sr^2^** | ***p*** | **Fit** |
| --- | --- | --- | --- | --- | --- | --- |
| (Intercept) | 0.28*** | [0.16, 0.40] |  |  | <.001 |  |
| Sex (Female) | -0.03** | [-0.04, -7.86 × 10^-3^] | -0.20 | 0.05 | .006 |  |
| Ponderal Index | 8.87 × 10^-3^* | [1.36 × 10^-3^, 0.02] | 0.16 | 0.03 | .02 |  |
| Maternal Age at Term | 2.51 × 10^-3^* | [5.37 × 10^-4^, 4.48 × 10^-3^] | 0.18 | 0.04 | .01 |  |
| Education (University) | 0.01 | [-4.03 × 10^-3^, 0.03] | 0.11 | 0.01 | .13 |  |
| CDS 14 gestational weeks | 2.56 × 10^-3^*** | [1.25 × 10^-3^, 3.86 × 10^-3^] | 0.27 | 0.08 | <.001 |  |
|  |  |  |  |  |  | R^2^ *=* .191*** |
|  |  |  |  |  |  | 95% CI [.08, .27] |

*** = p ≤ 0.001, ** = p ≤ 0.01, * = p ≤ 0.05

Note. ROI = Region of Interest. CDS = Composite Distress Score. A significant b-weight indicates the β-weight and semi-partial correlation are also significant. b represents unstandardized regression weights. β indicates the standardized regression weights. sr^2^ represents the semi-partial correlation squared. LL and UL indicate the lower and upper limits of a confidence interval, respectively. Confidence intervals are reported with sufficient decimal places to convey meaningful precision. Values with absolute values less than 0.01 are presented in scientific notation.

Table S5. Regression results using model 1b and cluster at 3 months ROI 1 (-6, 6, 60) as the criterion

| **Predictor** | **b** | **b**  **95% CI**  **[LL, UL]** | **β** | **sr^2^** | ***p*** | **Fit** |
| --- | --- | --- | --- | --- | --- | --- |
| (Intercept) | 0.72*** | [0.47, 0.96] |  |  | <.001 |  |
| Age At Scan | -1.41 × 10^-4^* | [-2.62 × 10^-4^, -1.91 × 10^-5^] | -0.15 | 0.02 | .02 |  |
| Sex (Female) | -0.03*** | [-0.04, -0.02] | -0.36 | 0.13 | <.001 |  |
| Maternal Age at Term | 1.01 × 10^-3^ | [-2.22 × 10^-4^, -1.37 × 10^-3^] | 0.11 | 0.01 | .11 |  |
| CDS 3 months | -2.54 × 10^-3^*** | [-3.71 × 10^-3^, -1.37 × 10^-3^] | -0.37 | 0.08 | <.001 |  |
| Prenatal CDS | 2.88 × 10^-4^ | [-1.08 × 10^-4^, 6.84 × 10^-4^] | 0.13 | 0.01 | .16 |  |
|  |  |  |  |  |  | R^2^ *=* .252*** |
|  |  |  |  |  |  | 95% CI [.13, .34] |

*** = p ≤ 0.001, ** = p ≤ 0.01, * = p ≤ 0.05

Note. ROI = Region of Interest. CDS = Composite Distress Score. A significant b-weight indicates the β-weight and semi-partial correlation are also significant. b represents unstandardized regression weights. β indicates the standardized regression weights. sr^2^ represents the semi-partial correlation squared. LL and UL indicate the lower and upper limits of a confidence interval, respectively. Confidence intervals are reported with sufficient decimal places to convey meaningful precision. Values with absolute values less than 0.01 are presented in scientific notation.

Table S6. Regression results using model 1b and 3 months ROI 2 (-8, -62, -33) as the criterion

| **Predictor** | **b** | **b**  **95% CI**  **[LL, UL]** | **β** | **sr^2^** | ***p*** | **Fit** |
| --- | --- | --- | --- | --- | --- | --- |
| (Intercept) | 0.46*** | [0.45, 0.47] |  |  | <.001 |  |
| Sex (Female) | -0.19*** | [-0.03, -7.63 × 10^-3^] | -0.23 | 0.06 | .001 |  |
| CDS 3 months | -1.79 × 10^-3^*** | [-2.70 × 10^-3^, -8.84 × 10^-4^] | -0.28 | 0.08 | <.001 |  |
|  |  |  |  |  |  | R^2^ = .124*** |
|  |  |  |  |  |  | 95% CI [.04, .21] |

*** = p ≤ 0.001, ** = p ≤ 0.01, * = p ≤ 0.05

Note. ROI = Region of Interest. CDS = Composite Distress Score. A significant b-weight indicates the β-weight and semi-partial correlation are also significant. b represents unstandardized regression weights. β indicates the standardized regression weights. sr^2^ represents the semi-partial correlation squared. LL and UL indicate the lower and upper limits of a confidence interval, respectively. Confidence intervals are reported with sufficient decimal places to convey meaningful precision. Values with absolute values less than 0.01 are presented in scientific notation.

Table S7. Regression results using model 1b and 3 months ROI 3 (-34, 8, 10) as the criterion

| **Predictor** | **b** | **b**  **95% CI**  **[LL, UL]** | **β** | **sr^2^** | ***p*** | **Fit** |
| --- | --- | --- | --- | --- | --- | --- |
| (Intercept) | 0.59*** | [0.58, 0.60] |  |  | <.001 |  |
| Sex (Female) | -0.05*** | [-0.06, -0.03] | -0.38 | 0.15 | <.001 |  |
| CDS 3 months | -2.25 × 10^-3^*** | [-3.49 × 10^-3^, -1.00 × 10^-3^] | -0.24 | 0.07 | <.001 |  |
|  |  |  |  |  |  | R^2^ *=* .196*** |
|  |  |  |  |  |  | 95% CI [.09, .29] |

*** = p ≤ 0.001, ** = p ≤ 0.01, * = p ≤ 0.05

Note. ROI = Region of Interest. CDS = Composite Distress Score. A significant b-weight indicates the β-weight and semi-partial correlation are also significant. b represents unstandardized regression weights. β indicates the standardized regression weights. sr^2^ represents the semi-partial correlation squared. LL and UL indicate the lower and upper limits of a confidence interval, respectively. Values with absolute values less than 0.01 are presented in scientific notation.

Table S8. Regression results using model 1b and 3 months ROI 4 (12, 44, 40) as the criterion

| **Predictor** | **b** | **b**  **95% CI**  **[LL, UL]** | **Β** | **sr^2^** | ***p*** | **Fit** |
| --- | --- | --- | --- | --- | --- | --- |
| (Intercept) | 0.38*** | [0.33, 0.43] |  |  | <.001 |  |
| Sex (Female) | -0.03*** | [-0.04, -0.01] | -0.25 | 0.07 | <.001 |  |
| Maternal Age at Term | 1.25 × 10^-3^ | [-3.26 × 10^-4^, 2.83 × 10^-3^] | 0.11 | 0.01 | .122 |  |
| Education (University) | 1.34 × 10^-3^ | [-1.43 × 10^-3^, 0.03] | 0.13 | 0.02 | .078 |  |
| CDS 3 months | -2.11 × 10^-3^*** | [-3.25 × 10^-3^, -9.68 × 10^-4^] | -0.26 | 0.07 | <.001 |  |
|  |  |  |  |  |  | R^2^ *=* .174*** |
|  |  |  |  |  |  | 95% CI [.07, .26] |

*** = p ≤ 0.001, ** = p ≤ 0.01, * = p ≤ 0.05

Note. ROI = Region of Interest. CDS = Composite Distress Score. A significant b-weight indicates the β-weight and semi-partial correlation are also significant. b represents unstandardized regression weights. β indicates the standardized regression weights. sr^2^ represents the semi-partial correlation squared. LL and UL indicate the lower and upper limits of a confidence interval, respectively. Confidence intervals are reported with sufficient decimal places to convey meaningful precision. Values with absolute values less than 0.01 are presented in scientific notation.

Table S9. Regression results using model 1b and 3 months ROI 5 (28, -64, -20) as the criterion

| **Predictor** | **b** | **b**  **95% CI**  **[LL, UL]** | **β** | **sr^2^** | ***p*** | **Fit** |
| --- | --- | --- | --- | --- | --- | --- |
| (Intercept) | 0.86** | [0.84, 0.88] |  |  | <.001 |  |
| Sex (Female) | -0.08** | [-0.10, -0.06] | -0.50 | 0.26 | <.001 |  |
| CDS 3 months | -4.09 × 10^-3^*** | [-6.24 × 10^-3^, -1.94 × 10^-3^] | -0.32 | 0.08 | <.001 |  |
| Prenatal CDS | 6.22 × 10^-4^ | [-1.02 × 10^-4^, 1.35 × 10^-3^] | 0.14 | 0.02 | .094 |  |
|  |  |  |  |  |  | R^2^ *=* .288*** |
|  |  |  |  |  |  | 95% CI [.17, .38] |

*** = p ≤ 0.001, ** = p ≤ 0.01, * = p ≤ 0.05

Note. ROI = Region of Interest. CDS = Composite Distress Score. A significant b-weight indicates the β-weight and semi-partial correlation are also significant. b represents unstandardized regression weights. β indicates the standardized regression weights. sr^2^ represents the semi-partial correlation squared. r represents the zero-order correlation. LL and UL indicate the lower and upper limits of a confidence interval, respectively. Confidence intervals are reported with sufficient decimal places to convey meaningful precision. Values with absolute values less than 0.01 are presented in scientific notation.

Table S10. Regression results using model 1b and 3 months ROI 6 (58, -28, 48) as the criterion

| **Predictor** | **b** | **b**  **95% CI**  **[LL, UL]** | **β** | **sr^2^** | ***p*** | **Fit** |
| --- | --- | --- | --- | --- | --- | --- |
| (Intercept) | 0.43*** | [0.42, 0.44] |  |  | <.001 |  |
| Sex (Female) | -0.02*** | [-0.04, -0.01] | -0.24 | 0.06 | .001 |  |
| CDS 3 months | -2.01 × 10^-3^*** | [-3.17 × 10^-3^, -8.57 × 10^-4^] | -0.25 | 0.06 | <.001 |  |
|  |  |  |  |  |  | R^2^ *=* .112*** |
|  |  |  |  |  |  | 95% CI [.03, .20] |

*** = p ≤ 0.001, ** = p ≤ 0.01, * = p ≤ 0.05

Note. ROI = Region of Interest. CDS = Composite Distress Score. A significant b-weight indicates the β-weight and semi-partial correlation are also significant. b represents unstandardized regression weights. β indicates the standardized regression weights. sr^2^ represents the semi-partial correlation squared. LL and UL indicate the lower and upper limits of a confidence interval, respectively. Confidence intervals are reported with sufficient decimal places to convey meaningful precision. Values with absolute values less than 0.01 are presented in scientific notation.

Table S11. Regression results using model 1b and 6 months ROI (10, -76, 4) as the criterion

| **Predictor** | **b** | **b**  **95% CI**  **[LL, UL]** | **β** | **sr^2^** | ***p*** | **Fit** |
| --- | --- | --- | --- | --- | --- | --- |
| (Intercept) | 0.49*** | [0.47, 0.51] |  |  | <.001 |  |
| Sex (Female) | -0.03** | [-0.04, -0.01] | -0.22 | 0.05 | .002 |  |
| Education (University) | -0.01 | [-0.03, 3.40 × 10^-3^] | -0.11 | 0.01 | .12 |  |
| CDS 6 months | 2.12 × 10^-3^*** | [1.05 × 10^-3^, 3.28 × 10^-3^] | 0.27 | 0.08 | <.001 |  |
|  |  |  |  |  |  | R^2^ = .127*** |
|  |  |  |  |  |  | 95% CI [.04, .21] |

*** = p ≤ 0.001, ** = p ≤ 0.01, * = p ≤ 0.05

Note. ROI = Region of Interest. CDS = Composite Distress Score. A significant b-weight indicates the β-weight and semi-partial correlation are also significant. b represents unstandardized regression weights. β indicates the standardized regression weights. sr^2^ represents the semi-partial correlation squared. LL and UL indicate the lower and upper limits of a confidence interval, respectively. Confidence intervals are reported with sufficient decimal places to convey meaningful precision. Values with absolute values less than 0.01 are presented in scientific notation.

Table S12. Regression results using model 1b and 24 months ROI 1 (-66, -48, 22) as the criterion

| **Predictor** | **b** | **b**  **95% CI**  **[LL, UL]** | **β** | **sr^2^** | ***p*** | **Fit** |
| --- | --- | --- | --- | --- | --- | --- |
| (Intercept) | 0.37*** | [0.25, 0.50] |  |  | <.001 |  |
| Sex (Female) | -0.04*** | [-0.06, -0.02] | -0.28 | 0.08 | <.001 |  |
| Ponderal Index | 0.01* | [1.36 × 10^-3^, 0.02] | 0.16 | 0.03 | .02 |  |
| CDS 2 years | -2.50 × 10^-3^*** | [-3.98 × 10^-3^, -1.02 × 10^-3^] | -0.24 | 0.06 | .001 |  |
|  |  |  |  |  |  | R^2^ = .142*** |
|  |  |  |  |  |  | 95% CI [.05, .23] |

*** = p ≤ 0.001, ** = p ≤ 0.01, * = p ≤ 0.05

Note. ROI = Region of Interest. CDS = Composite Distress Score. A significant b-weight indicates the β-weight and semi-partial correlation are also significant. b represents unstandardized regression weights. β indicates the standardized regression weights. sr^2^ represents the semi-partial correlation squared. LL and UL indicate the lower and upper limits of a confidence interval, respectively. Confidence intervals are reported with sufficient decimal places to convey meaningful precision. Values with absolute values less than 0.01 are presented in scientific notation.

Table S13. Regression results using model 1b and 24 months ROI 2 (14, 54, 24) as the criterion

| **Predictor** | **b** | **b**  **95% CI**  **[LL, UL]** | **β** | **sr^2^** | ***p*** | **Fit** |
| --- | --- | --- | --- | --- | --- | --- |
| (Intercept) | 0.42*** | [0.32, 0.51] |  |  | <.001 |  |
| Sex (Female) | -0.03*** | [-0.04, -0.01] | -0.28 | 0.09 | <.001 |  |
| Ponderal Index | -4.20 × 10^-3^ | [-0.01, 1.66 × 10^-3^] | -0.10 | 0.01 | .162 |  |
| Maternal Age at Term | 2.02 × 10^-3^** | [4.92 × 10^-4^, 3.54 × 10^-3^] | 0.18 | 0.04 | .01 |  |
| Education (University) | 0.01 | [-1.91 × 10^-3^, 0.03] | 0.12 | 0.02 | .091 |  |
| CDS 2 years | -1.87 × 10^-3^*** | [-2.89 × 10^-3^, -8.46 × 10^-4^] | -0.25 | 0.07 | <.001 |  |
|  |  |  |  |  |  | R^2^ = .211*** |
|  |  |  |  |  |  | 95% CI [.09, .29] |

*** = p ≤ 0.001, ** = p ≤ 0.01, * = p ≤ 0.05

Note. ROI = Region of Interest. CDS = Composite Distress Score. A significant b-weight indicates the β-weight and semi-partial correlation are also significant. b represents unstandardized regression weights. β indicates the standardized regression weights. sr^2^ represents the semi-partial correlation squared. LL and UL indicate the lower and upper limits of a confidence interval, respectively. Confidence intervals are reported with sufficient decimal places to convey meaningful precision. Values with absolute values less than 0.01 are presented in scientific notation.

Table S14. Summary of models 2 fit statistics for ROI eigenvalues ~ CDS from the time point in which the ROI was significant + Age at scan + Sex + Ponderal Index + Maternal age at term + Maternal education + Prenatal alcohol exposure + Prenatal smoking exposure + Prenatal selective serotonin reuptake inhibitor or serotonin and norepinephrine reuptake inhibitor exposure + Obstetric risk + Maternal pre-pregnancy body mass index + Neonatal intensive care unit admission

|  | **F (df1, df2)** | ***p*** | **R^2^** | **Adjusted R^2^** |
| --- | --- | --- | --- | --- |
| **14 gestational weeks** Right precuneus/ superior parietal lobule (12, -72, 50) | 7.91*** (5, 167) | <.001 | .19 | .17 |
| **3 months (ROI 1)** Left supplementary motor cortex  (-6, 6, 60) | 13.52*** (4, 168) | <.001 | .24 | .23 |
| **3 months (ROI 2)** Cerebellar vermal lobules VIII-X/left cerebellum  (-8, -62, -33) | 7.33*** (4, 168) | <.001 | .15 | .13 |
| **3 months (ROI 3)** Left anterior insula  (-34, 8, 10) | 12.20*** (4, 168) | <.001 | .23 | .21 |
| **3 months (ROI 4)** Right superior frontal gyrus  (12, 44, 40) | 9.00*** (4, 168) | <.001 | .18 | .16 |
| **3 months (ROI 5)** Right cerebellum (28, -64, -20) | 17.88*** (4, 168) | <.001 | .30 | .28 |
| **3 months (ROI 6)** Right supramarginal gyrus  (58, -28, 48) | 7.51*** (4, 168) | <.001 | .15 | .13 |
| **6 months**  Right calcarine cortex/ lingual gyrus (10, -76, 4) | 5.94*** (5, 167) | <.001 | .15 | .13 |
| **2 years (ROI 1)** Left supramarginal gyrus  (-66, -48, 22) | 7.12*** (7, 165) | <.001 | .23 | .20 |
| **2 years (ROI 2)** Right superior frontal gyrus (14, 54, 24) | 7.19*** (5, 167) | <.001 | .18 | .15 |

*** = p ≤ 0.001, ** = p ≤ 0.01, * = p ≤ 0.05

CDS = Composite Distress Score, () = coordinates (x, y, z), ROI = region of interest

Table S15. Regression results using model 2 and 14 gestational weeks ROI (12, -72, 50) as the criterion

| **Predictor** | **b** | **b**  **95% CI**  **[LL, UL]** | **β** | **sr^2^** | ***p*** | **Fit** |
| --- | --- | --- | --- | --- | --- | --- |
| (Intercept) | 0.28*** | [0.16, 0.40] |  |  | <.001 |  |
| CDS 14 gestational weeks | 2.56 × 10^-3^*** | [1.25 × 10^-3^, 3.86 × 10^-3^] | 0.27 | 0.07 | <.001 |  |
| Sex (Female) | -0.03** | [-0.04, -7.86 × 10^-3^] | -0.20 | 0.04 | .006 |  |
| Ponderal Index | 0.01* | [1.36 × 10^-3^, 0.02] | 0.16 | 0.03 | .02 |  |
| Maternal Age at Term | 2.51 × 10^-3^** | [5.37 × 10^-4^, 4.48 × 10^-3^] | 0.18 | 0.03 | .01 |  |
| Education (University) | 0.01 | [-4.03 × 10^-3^, 0.03] | 0.11 | 0.01 | .128 |  |
|  |  |  |  |  |  | R^2^ = .191*** |
|  |  |  |  |  |  | 95% CI [.08, .27] |

*** = p ≤ 0.001, ** = p ≤ 0.01, * = p ≤ 0.05

Note. ROI = Region of Interest. CDS = Composite Distress Score. A significant b-weight indicates the β-weight and semi-partial correlation are also significant. b represents unstandardized regression weights. β indicates the standardized regression weights. sr^2^ represents the semi-partial correlation squared. LL and UL indicate the lower and upper limits of a confidence interval, respectively. Confidence intervals are reported with sufficient decimal places to convey meaningful precision. Values with absolute values less than 0.01 are presented in scientific notation.

Table S16. Regression results using model 2 and 3 months ROI 1 (-6, 6, 60) as the criterion

| **Predictor** | **b** | **b**  **95% CI**  **[LL, UL]** | **β** | **sr^2^** | ***p*** | **Fit** |
| --- | --- | --- | --- | --- | --- | --- |
| (Intercept) | 0.78*** | [0.54, 1.02] |  |  | <.001 |  |
| CDS 3 months | -1.96 × 10^-3^*** | [-2.87 × 10^-3^, -1.06 × 10^-3^] | -0.29 | 0.10 | <.001 |  |
| Age at Scan | -1.54 × 10^-4^ ** | [-2.74 × 10^-4^, -3.30 × 10^-5^] | -0.17 | 0.04 | .01 |  |
| Sex (Female) | -0.03*** | [-0.04, -0.02] | -0.37 | 0.15 | <.001 |  |
| Smoking (Yes) | -0.02 | [-0.04, 3.95 × 10^-3^] | -0.11 | 0.02 | 0.10 |  |
|  |  |  |  |  |  | *R^2^* = .244*** |
|  |  |  |  |  |  | 95% CI [.13, .33] |

*** = p ≤ 0.001, ** = p ≤ 0.01, * = p ≤ 0.05

Note. ROI = Region of Interest. CDS = Composite Distress Score. A significant b-weight indicates the β-weight and semi-partial correlation are also significant. b represents unstandardized regression weights. β indicates the standardized regression weights. sr^2^ represents the semi-partial correlation squared. LL and UL indicate the lower and upper limits of a confidence interval, respectively. Confidence intervals are reported with sufficient decimal places to convey meaningful precision. Values with absolute values less than 0.01 are presented in scientific notation.

Table S17. Regression results using model 2 and 3 months ROI 2 (-8, -62, -33) as the criterion

| **Predictor** | **b** | **b**  **95% CI**  **[LL, UL]** | **β** | **sr^2^** | ***p*** | **Fit** |
| --- | --- | --- | --- | --- | --- | --- |
| (Intercept) | 0.49*** | [0.45, 0.54] |  |  | <.001 |  |
| CDS 3 months | -1.75 × 10^-3^ *** | [-2.66 × 10^-3^, -8.43 × 10^-4^] | -0.27 | 0.08 | <.001 |  |
| Sex (Female) | -0.02*** | [-0.03, -8.79 × 10^-3^] | -0.25 | 0.07 | <.001 |  |
| Maternal Age at Term | -1.07 × 10^-3^ | [-2.33 × 10^-3^, 1.88 × 10^-4^] | -0.12 | 0.02 | .10 |  |
| Smoking (Yes) | -0.02 | [-0.05, 2.54 × 10^-3^] | -0.13 | 0.02 | .08 |  |
|  |  |  |  |  |  | *R^2^* = .149*** |
|  |  |  |  |  |  | 95% CI [.05, .23] |

*** = p ≤ 0.001, ** = p ≤ 0.01, * = p ≤ 0.05

Note. ROI = Region of Interest. CDS = Composite Distress Score. A significant b-weight indicates the β-weight and semi-partial correlation are also significant. b represents unstandardized regression weights. β indicates the standardized regression weights. sr^2^ represents the semi-partial correlation squared. LL and UL indicate the lower and upper limits of a confidence interval, respectively. Confidence intervals are reported with sufficient decimal places to convey meaningful precision. Values with absolute values less than 0.01 are presented in scientific notation.

Table S18. Regression results using model 2 and 3 months ROI 3 (-34, 8, 10) as the criterion

| **Predictor** | **b** | **b**  **95% CI**  **[LL, UL]** | **β** | **sr^2^** | ***p*** | **Fit** |
| --- | --- | --- | --- | --- | --- | --- |
| (Intercept) | 0.59*** | [0.57, 0.60] |  |  | <.001 |  |
| CDS 3 months | -2.12 × 10^-3^ *** | [-3.36 × 10^-3^, -8.81 × 10^-4^] | -0.23 | 0.06 | <.001 |  |
| Sex (Female) | -0.04*** | [-0.06, -0.03] | -0.37 | 0.15 | <.001 |  |
| Smoking (Yes) | -0.03 | [-0.06, 3.57 × 10^-3^] | -0.12 | 0.02 | .08 |  |
| NICU (Yes) | 0.02* | [2.22 × 10^-4^, 0.04] | 0.14 | 0.02 | .05 |  |
|  |  |  |  |  |  | *R^2^* = .225*** |
|  |  |  |  |  |  | 95% CI [.11, .31] |

*** = p ≤ 0.001, ** = p ≤ 0.01, * = p ≤ 0.05

Note. ROI = Region of Interest. CDS = Composite Distress Score. A significant b-weight indicates the β-weight and semi-partial correlation are also significant. b represents unstandardized regression weights. β indicates the standardized regression weights. sr^2^ represents the semi-partial correlation squared. LL and UL indicate the lower and upper limits of a confidence interval, respectively. Confidence intervals are reported with sufficient decimal places to convey meaningful precision. Values with absolute values less than 0.01 are presented in scientific notation.

Table S19. Regression results using model 2 and 3 months ROI 4 (12, 44, 40) as the criterion

| **Predictor** | **b** | **b**  **95% CI**  **[LL, UL]** | **β** | **sr^2^** | ***p*** | **Fit** |
| --- | --- | --- | --- | --- | --- | --- |
| (Intercept) | 0.42*** | [0.40, 0.43] |  |  | <.001 |  |
| CDS 3 months | -2.22 × 10^-3^ *** | [-3.35 × 10^-3^, -1.09 × 10^-3^] | -.27 | 0.08 | <.001 |  |
| Sex (Female) | -0.03*** | [-0.04, -0.01] | -.25 | 0.07 | <.001 |  |
| Education (University) | 0.02* | [1.02 × 10^-3^, 0.03] | .25 | 0.03 | .04 |  |
| NICU (Yes) | 0.02 | [-2.66 × 10^-3^, 0.04] | .12 | 0.02 | .09 |  |
|  |  |  |  |  |  | *R^2^* = .176*** |
|  |  |  |  |  |  | 95% CI [.07, .26] |

*** = p ≤ 0.001, ** = p ≤ 0.01, * = p ≤ 0.05

Note. ROI = Region of Interest. CDS = Composite Distress Score. A significant b-weight indicates the β-weight and semi-partial correlation are also significant. b represents unstandardized regression weights. β indicates the standardized regression weights. sr^2^ represents the semi-partial correlation squared. LL and UL indicate the lower and upper limits of a confidence interval, respectively. Confidence intervals are reported with sufficient decimal places to convey meaningful precision. Values with absolute values less than 0.01 are presented in scientific notation.

Table S20. Regression results using model 2 and 3 months ROI 5 (28, -64, -20) as the criterion

| **Predictor** | **b** | **b**  **95% CI**  **[LL, UL]** | **β** | **sr^2^** | ***p*** | **Fit** |
| --- | --- | --- | --- | --- | --- | --- |
| (Intercept) | 0.92*** | [0.85, 1.00] |  |  | <.001 |  |
| CDS 3 months | -2.80 × 10^-3^ *** | [-0.004, -0.001] | -.22 | 0.06 | .001 |  |
| Sex (Female) | -0.08*** | [-0.10, -0.06] | -.50 | 0.26 | <.001 |  |
| Maternal Age at Term | -1.76 × 10^-3^ | [-0.004, 5.40 × 10^-4^] | -.10 | 0.01 | .14 |  |
| Smoking (Yes) | -0.05* | [-0.09, -0.001] | -.14 | 0.02 | .04 |  |
|  |  |  |  |  |  | *R^2^* = .299*** |
|  |  |  |  |  |  | 95% CI [.18, .39] |

*** = p ≤ 0.001, ** = p ≤ 0.01, * = p ≤ 0.05

Note. ROI = Region of Interest. CDS = Composite Distress Score. A significant b-weight indicates the β-weight and semi-partial correlation are also significant. b represents unstandardized regression weights. β indicates the standardized regression weights. sr^2^ represents the semi-partial correlation squared. LL and UL indicate the lower and upper limits of a confidence interval, respectively. Confidence intervals are reported with sufficient decimal places to convey meaningful precision. Values with absolute values less than 0.01 are presented in scientific notation.

Table S21. Regression results using model 2 and 3 months ROI 6 (58, -28, 48) as the criterion

| **Predictor** | **b** | **b**  **95% CI**  **[LL, UL]** | **β** | **sr^2^** | ***p*** | **Fit** |
| --- | --- | --- | --- | --- | --- | --- |
| (Intercept) | 0.39*** | [0.33, 0.44] |  |  | <.001 |  |
| CDS 3 months | -1.84 × 10^-3^ ** | [-2.98 × 10^-3^, -7.03 × 10^-4^] | -.23 | 0.06 | .002 |  |
| Sex (Female) | -0.02** | [-0.04, -8.96 × 10^-3^] | -.23 | 0.06 | .002 |  |
| Maternal Age at Term | 1.48 × 10^-3^ | [-1.02 × 10^-4^, 3.06 × 10^-3^] | .13 | 0.02 | .07 |  |
| Obstetric risk (Yes) | -0.03* | [-0.05, -5.12 × 10^-3^] | -.18 | 0.03 | .02 |  |
|  |  |  |  |  |  | *R^2^* = .152*** |
|  |  |  |  |  |  | 95% CI [.05, .23] |

*** = p ≤ 0.001, ** = p ≤ 0.01, * = p ≤ 0.05

Note. ROI = Region of Interest. CDS = Composite Distress Score. A significant b-weight indicates the β-weight and semi-partial correlation are also significant. b represents unstandardized regression weights. β indicates the standardized regression weights. sr^2^ represents the semi-partial correlation squared. LL and UL indicate the lower and upper limits of a confidence interval, respectively. Confidence intervals are reported with sufficient decimal places to convey meaningful precision. Values with absolute values less than 0.01 are presented in scientific notation.

Table S22. Regression results using model 2 and 6 months ROI (10, -76, 4) as the criterion

| **Predictor** | **b** | **b**  **95% CI**  **[LL, UL]** | **β** | **sr^2^** | ***p*** | **Fit** |
| --- | --- | --- | --- | --- | --- | --- |
| (Intercept) | 0.49*** | [0.47, 0.51] |  |  | <.001 |  |
| CDS 6 months | 2.18 × 10^-3^ *** | [1.07 × 10^-3^, 3.29 × 10^-3^] | .28 | 0.08 | <.001 |  |
| Sex (Female) | -0.03** | [-0.04, -8.44 × 10^-3^] | -.21 | 0.05 | .004 |  |
| Education (University) | -0.02 | [-0.03, 8.70 × 10^-4^] | -.14 | 0.02 | .06 |  |
| Smoking (Yes) | -0.03 | [-0.06, 3.93 × 10^-3^] | -.13 | 0.02 | .08 |  |
| NICU (Yes) | 0.02 | [-5.94 × 10^-3^, 0.04] | .11 | 0.01 | .15 |  |
|  |  |  |  |  |  | *R^2^* = .151*** |
|  |  |  |  |  |  | 95% CI [.05, .23] |

*** = p ≤ 0.001, ** = p ≤ 0.01, * = p ≤ 0.05

Note. ROI = Region of Interest. CDS = Composite Distress Score. A significant b-weight indicates the β-weight and semi-partial correlation are also significant. b represents unstandardized regression weights. β indicates the standardized regression weights. sr^2^ represents the semi-partial correlation squared. LL and UL indicate the lower and upper limits of a confidence interval, respectively. Confidence intervals are reported with sufficient decimal places to convey meaningful precision. Values with absolute values less than 0.01 are presented in scientific notation.

Table S23. Regression results using model 2 and 24 months ROI 1 (-66, -48, 22) as the criterion

| **Predictor** | **b** | **b**  **95% CI**  **[LL, UL]** | **β** | **sr^2^** | ***p*** | **Fit** |
| --- | --- | --- | --- | --- | --- | --- |
| (Intercept) | 0.39*** | [0.27, 0.51] |  |  | <.001 |  |
| CDS 2 years | -2.26 × 10^-3^ ** | [-3.73 × 10^-3^, -7.92 × 10^-4^] | -.22 | 0.05 | .003 |  |
| Sex (Female) | -0.04*** | [-0.06, -0.02] | -.28 | 0.08 | <.001 |  |
| Ponderal Index | 9.33 × 10^-3^* | [9.17 × 10^-4^, 0.02] | .15 | 0.03 | .03 |  |
| Smoking (Yes) | -0.04* | [-0.08, -2.91 × 10^-3^] | -.15 | 0.03 | .04 |  |
| Obstetric risk (Yes) | -0.03 | [-0.05, 2.77 × 10^-3^] | -.12 | 0.02 | .08 |  |
|  |  |  |  |  |  | R2 = .177*** |
|  |  |  |  |  |  | 95% CI [.07, .26] |

*** = p ≤ 0.001, ** = p ≤ 0.01, * = p ≤ 0.05

Note. ROI = Region of Interest. CDS = Composite Distress Score. A significant b-weight indicates the β-weight and semi-partial correlation are also significant. b represents unstandardized regression weights. β indicates the standardized regression weights. sr^2^ represents the semi-partial correlation squared. LL and UL indicate the lower and upper limits of a confidence interval, respectively. Confidence intervals are reported with sufficient decimal places to convey meaningful precision. Values with absolute values less than 0.01 are presented in scientific notation.

Table S24. Regression results using model 2 and 24 months ROI 2 (14, 54, 24) as the criterion

| **Predictor** | **b** | **b**  **95% CI**  **[LL, UL]** | **β** | **sr^2^** | ***p*** | **Fit** |
| --- | --- | --- | --- | --- | --- | --- |
| (Intercept) | 0.43*** | [0.33, 0.52] |  |  | <.001 |  |
| CDS 2 years | -1.79 × 10^-3^*** | [-2.81 × 10^-3^, -7.65 × 10^-4^] | -.24 | .07 | <.001 |  |
| Sex (Female) | -0.03*** | [-0.04, -0.01] | -.27 | .08 | <.001 |  |
| Ponderal Index | -4.17 × 10^-3^ | [-9.98 × 10^-3^, 1.65 × 10^-3^] | -.10 | .01 | .16 |  |
| Maternal Age at Term | 1.66 × 10^-3^* | [1.14 × 10^-4^, 3.22 × 10^-3^] | .15 | .03 | .04 |  |
| Education (University) | 0.01 | [-3.43 × 10^-3^, 0.03] | .11 | .01 | .14 |  |
| Smoking (Yes) | -0.02 | [-0.05, 6.42 × 10^-3^] | -.11 | .01 | .13 |  |
| NICU (Yes) | 0.02 | [-3.01 × 10^-3^, 0.04] | .12 | .02 | .10 |  |
|  |  |  |  |  |  | *R^2^* = .232*** |
|  |  |  |  |  |  | 95% CI [.10, .31] |

*** = p ≤ 0.001, ** = p ≤ 0.01, * = p ≤ 0.05

Note. ROI = Region of Interest. CDS = Composite Distress Score. A significant b-weight indicates the β-weight and semi-partial correlation are also significant. b represents unstandardized regression weights. β indicates the standardized regression weights. sr^2^ represents the semi-partial correlation squared. LL and UL indicate the lower and upper limits of a confidence interval, respectively. Confidence intervals are reported with sufficient decimal places to convey meaningful precision. Values with absolute values less than 0.01 are presented in scientific notation.

##### Controlling for other composite distress score timepoints

Compared to the main models, 14 GW results disappeared, while 34 GW results are completely new. Results at 3 months postpartum are all negative, like in the main models. Out of the significant clusters, three out of six (the left supplementary motor cortex, left anterior insula, and right cerebellum / right fusiform gyrus) are similar to those in the main models, while the other three are in different regions than the results form the main model. At 6 months postpartum, the medial occipital lobe cluster (including regions such as bilateral calcarine and lingual cortex) reflects the result from the main model. In addition, four new significant clusters appeared. All of them were positive. The 24 months postpartum was the only time point in which the direction of the results changed (negative in the main models, positive in sensitivity analyses). Notably, the sensitivity results appear bilaterally in the posterior caudate nucleus, extending to surrounding structures such as the lateral ventricles, therefore to a large extent representing gray-white matter boundary effects related to the use of unthresholded smoothed gray matter probability maps. The information of the significant clusters is presented in Table S25 and the results are visualized in Figure S4.

The supplementary analyses are similar to the main analyses regarding the prenatal to postnatal ratio of the results. Multiple significant clusters associated with postnatal distress, and relatively fewer results associated with prenatal distress.

Table S25. Clusters from the supplementary voxel-based morphometry analysis.

| Region | x | y | z | Cluster size | p (FDR-corrected) | Peak z |
| --- | --- | --- | --- | --- | --- | --- |
| **34 gestational weeks (positive association)** | | | | | | |
| Right inferior occipital gyrus | 33 | -86 | 6 | 509 | 0.035 | 3.91 |
| Right posterior insula (gray−white matter boundary effect) | 28 | -24 | 4 | 875 | 0.013 | 3.86 |
| Right superior occipital gyrus | 26 | -80 | 20 | 518 | 0.035 | 3.85 |
| Left cerebellum | -12 | -86 | -38 | 545 | 0.035 | 3.08 |
| **3 months postpartum (negative association)** | | | | | | |
| Right cerebellum / right occipital fusiform gyrus | 33 | -66 | -20 | 1521 | < 0.001 | 4.03 |
| Left supplementary motor cortex | -4 | 8 | 58 | 1279 | 0.001 | 3.95 |
| Left anterior insula | -38 | 10 | 3 | 1671 | < 0.001 | 3.72 |
| Left medial/posterior orbital gyrus | -20 | 10 | -24 | 841 | 0.008 | 3.70 |
| Left cerebellum | -33 | -52 | -40 | 718 | 0.011 | 3.41 |
| Left inferior temporal gyrus | -52 | -4 | -44 | 770 | 0.010 | 3.37 |
| **6 months postpartum (positive association)** | | | | | | |
| Bilateral cuneus/calcarine cortex/lingual gyrus | -21 | -86 | 0 | 3768 | < 0.001 | 3.98 |
| Right inferior occipital gyrus | 44 | -66 | 9 | 682 | 0.025 | 3.97 |
| Right fusiform gyrus | 39 | -48 | -14 | 562 | 0.027 | 3.74 |
| Right precentral gyrus | 39 | -6 | 40 | 553 | 0.027 | 3.58 |
| Right insula | 45 | -3 | 2 | 533 | 0.027 | 3.51 |
| **24 months postpartum (positive association)** | | | | | | |
| Left caudate nucleus (gray−white matter boundary effect) | -18 | -26 | 22 | 638 | 0.022 | 3.75 |
| Right caudate nucleus (gray−white matter boundary effect) | 18 | -18 | 22 | 471 | 0.035 | 3.51 |

*Abbreviations: FDR = false discovery rate*

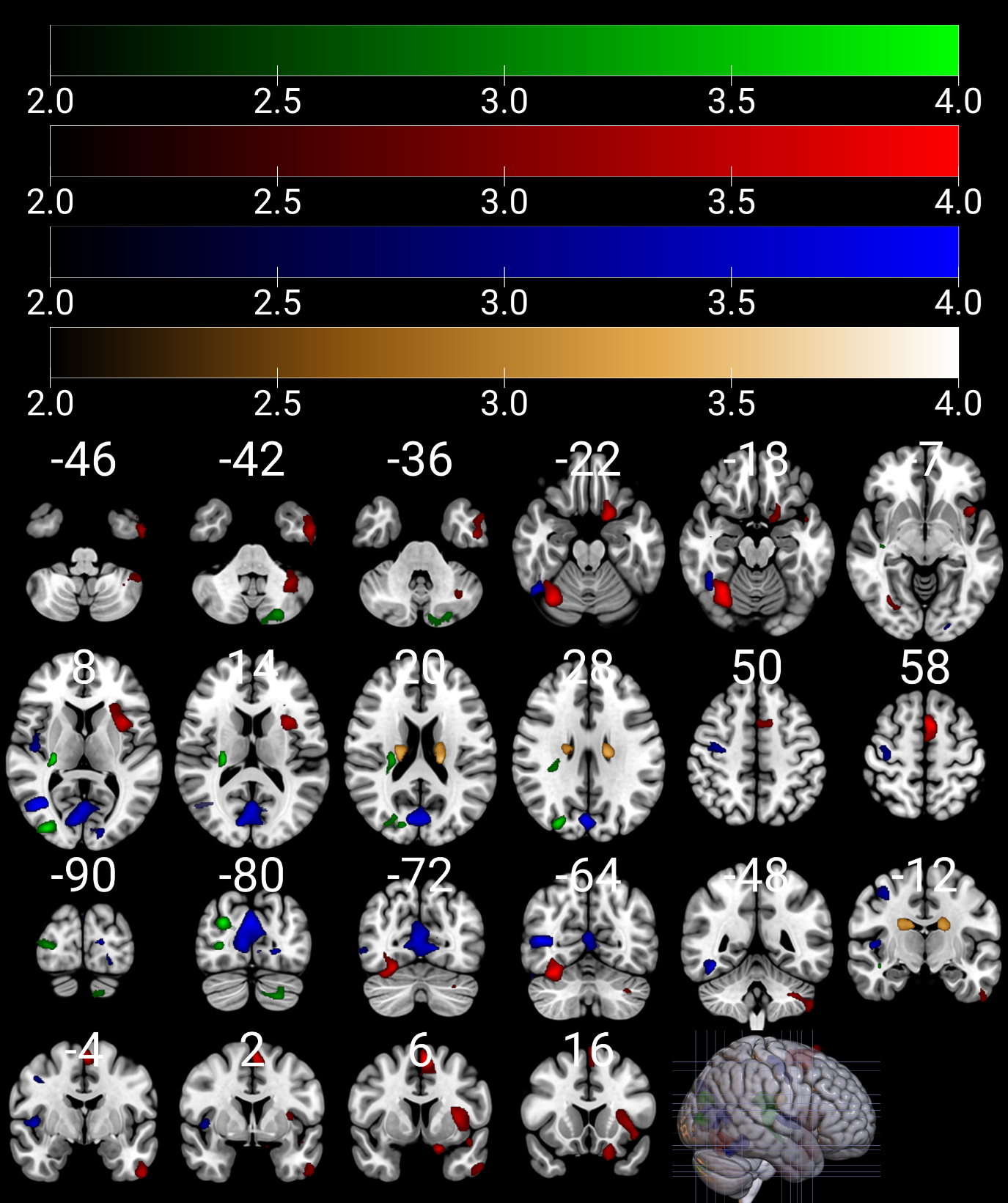

Figure S4. Color coding: Green = positive association between brain volume and composite distress score (CDS) at gestational week 34; Red = negative association between brain volume and composite distress score (CDS) at 3 months postnatal; Blue = positive association between brain volume and composite distress score (CDS) at 6 months postnatal; Gold = positive association between brain volume and composite distress score (CDS) at 24 months postnatal. Only statistically significant regions at p < 0.005, false discovery rate corrected at cluster level are displayed. Color indicates t-value. Images are in radiological orientation. Coordinates (indicated with white numbers) are in MNI space. The figure was created with MRIcroGL.
